# DeepNull: Modeling non-linear covariate effects improves phenotype prediction and association power

**DOI:** 10.1101/2021.05.26.445783

**Authors:** Farhad Hormozdiari, Zachary R. McCaw, Thomas Colthurst, Taedong Yun, Nick Furlotte, Andrew Carroll, Babak Alipanahi, Cory Y. McLean

## Abstract

Genome-wide association studies (GWAS) are among the workhorses of statistical genetics, having detected thousands of variants associated with complex traits and diseases. A typical GWAS examines the association between genotypes and the phenotype of interest while adjusting for a set of covariates. While covariates potentially have non-linear effects on the phenotype in many real world settings, due to the challenge of specifying the model, GWAS seldom include non-linear terms. Here we introduce DeepNull, a method that models non-linear covariate effects on phenotypes using a deep neural network (DNN) and then includes the model prediction as a single extra term in the GWAS association. First, using simulated data, we show that DeepNull increases statistical power by up to 20% while maintaining tight control of the type I error in the presence of interactions or non-linear covariate effects. Second, DeepNull maintains similar results to a standard GWAS when covariates have only linear effects on the phenotype. Third, DeepNull detects larger numbers of significant hits and loci (7% additional loci averaged over 10 traits) than standard GWAS in ten phenotypes from the UK Biobank (*n*=370K). Many of the hits found only by DeepNull are biologically plausible or have previously been reported in the GWAS catalog. Finally, DeepNull improves phenotype prediction by 23% averaged over the same ten phenotypes, the highest improvement was observed in the case of Glaucoma referral probability where DeepNull improves the phenotype prediction by 83%.

## 1 Introduction

GWAS aim to detect genetic variants or single-nucleotide polymorphisms (SNPs) that are associated with complex traits or diseases. Over the past decade, GWAS have successfully identified thousands of variants associated with common traits and disease [2, 4, 17, 20, 27, 39]. These associations have expanded our knowledge of biological mechanisms [7] and improved phenotype risk prediction [23].

In most GWAS, the association strength between genotype and phenotype is assessed while adjusting for a set of covariates, such as age, sex, and principal components (PCs) of the genetic relatedness matrix. Covariates are included in GWAS for two main reasons: to increase precision and to reduce confounding. In the linear model setting, adjustment for a covariate will improve precision if the distribution of the phenotype differs across levels of the covariate. For example, when performing GWAS on height, males and females have different means. Adjusting for sex reduces residual variation, and thereby increases power to detect an association between height and the candidate SNPs. Note, however, that omitting sex from the association test is entirely valid. In contrast, omitting a confounder will result in a biased test of association. By definition, a confounder is a common cause of the exposure (i.e. genotype) and the outcome (i.e. phenotype) [19]. An example confounder in GWAS is genetic ancestry: two ancestry groups may differ with respect to minor allele frequency (MAF) at common SNPs and, for unrelated reasons, in their phenotypic means. Failure to adjust for ancestry will lead to spurious associations between the phenotype and the SNPs whose MAFs differ, inflating the type I error of the association test. To reduce confounding due to population substructure, or the presence of genetically related subgroups within the cohort, multiple genetic PCs are commonly included as covariates during association testing [32, 37].

While most GWAS that adjust for covariates only include linear adjustments, recent GWAS have begun to model quadratic or first-order interaction terms between covariates [6, 26, 42]. For example, Shrine *et al*. [42] included `age`^2^ as a covariate when studying chronic obstructive pulmonary disease; Chen *et al*. [6] included squared body mass index (`BMI`^2^) when studying obstructive sleep apnea; and Kosmicki *et al*. [26] included an age by sex interaction (`age`×`sex`) when studying COVID-19 disease outcomes. Although these recent works have recognized the potential importance of modeling non-linear covariate effects, no systematic approach has been described for detecting the appropriate non-linear functions to adjust for in GWAS. The difficulty stems from the exponential number of possible interactions that can arise from a finite set of covariates (e.g. `age`^2^, `age` × `sex`, `age`^2^ × `sex`, …), and the infinite number of possible transformations of any given continuous covariate. Lastly, the optimal number of covariate interactions is not known *a priori* and requires testing different possible covariate interactions (Table S1).

In this work, we address the issue of GWAS model misspecification owing to non-linear influences of covariates on the phenotype of interest. We propose DeepNull, a method that enables modeling arbitrarily complex non-linearities between covariates and the phenotype of interest in GWAS associations in an unbiased manner. In simulated data, we show that DeepNull markedly improves association power and phenotype prediction in the presence of non-linear covariate effects on the phenotype, and retains equivalent performance in the absence of non-linear effects. We then demonstrate improvements in association power and phenotype prediction across ten phenotypes in the UK Biobank (UKB) [5], indicating DeepNull’s potential for broad utility in biobank-scale GWAS. We provide DeepNull as freely available open-source software (see URLs) for straightforward integration into existing GWAS association platforms.

## 2 Results

### 2.1 DeepNull overview

DeepNull trains a DNN to predict a phenotype of interest from covariates not directly derived from genotype data (hereafter “non-genetic covariates”). The DNN prediction can capture any possible complex non-linear relationship between the covariates and the phenotype. The DNN prediction for each individual in the GWAS cohort is then included as a single additional covariate within the standard GWAS model when performing genetic association testing. Adjusting for the DNN phenotype prediction in the final association model is equivalent to regressing it out of both the phenotype and the genotype. DeepNull reduces residual variation, thereby increasing statistical power (Figure S1).

Consider a quantitative phenotype ascertained for a sample of *n* individuals genotyped at *m* SNPs. Let 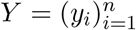 denote the *n* × 1 phenotype vector, where *y*_*i*_ is the phenotype value of the *i*-th individual; let ***G*** = [*g*_*ij*_] denote the *n* × *m* sample by SNP genotypes matrix, where *g*_*ij*_ is the minor allele count for the *i*-th individual at the *j*-th variant. Let 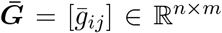denote the standardized version of ***G***, in which columns have been centered and scaled to have mean zero and unit variance. Furthermore, let *h* be a (possibly non-linear) function that predicts the phenotype from non-genetic covariates; we learn *h* using a DNN trained with cross-validation on the sample. The DeepNull association model is as follows:

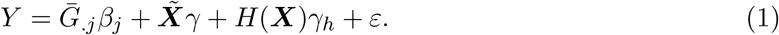

Here *β* is the *m* × 1 vector of effect sizes for each variant on the phenotype; 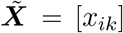 is the *n* × (*p* + *g*) covariate matrix that includes *p* non-genetic prognostic factors (*e*.*g*. age and sex) and *g* adjustments for genetic confounding (*e*.*g*. genetic PCs); *γ* is the (*p* + *g*) × 1 vector of association coefficients for all covariates. Compared to the standard GWAS association model, the DeepNull association model differs only by the inclusion of a single extra term *H*(***X***)*γ*_*h*_: ***X*** is the *n* × *p* subset of 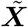 that excludes genetic confounders (see Methods); *H*: ℝ^*n*×*p*^ *→* ℝ^*n*^ is the function that applies *h* to each row of ***X***; *γ*_*h*_ is the scalar association coefficient for the DNN prediction of phenotype from non-genetic covariates.

### 2.2 DeepNull and Baseline model achieve similar results under linear model

We simulated phenotypes using UK Biobank [5] genotype data and used standardized `age`, `sex`, and `genotyping_array` as true covariates for 10,000 randomly selected individuals (Methods). First, we considered a linear effect for covariates on phenotypes (*f* (*x*) = *γx*). We simulated 100 phenotypes for each of six different genetic architectures with varying amounts of phenotypic variance explained by genetic data 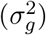 and by covariates 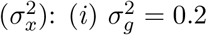 and 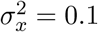, (*ii*) 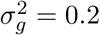 and 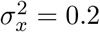, (*iii*) 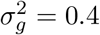 and 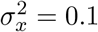, (*iv*) 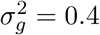 and 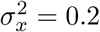, (*v*) 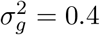 and 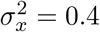, and (*vi*) 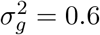 and 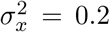. Causal variants were randomly embedded within `chr22` and non-causal variants within `chr1` and `chr2`. We compared DeepNull GWAS with standard GWAS (hereafter referred to as “Baseline”), each of which was performed using BOLT-LMM [30] (Methods). Statistical power and expected Chi-square statistics for the causal chromosome (`chr22`) are similar for DeepNull and Baseline (Figure 1a,b, Table S2). Statistical power for both DeepNull and Baseline increases as genetic heritability 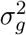 increases, which is expected since statistical power depends on both the effect size and genetic heritability. Additionally, the type I error is controlled and expected Chi-square statistics for non-causal variants are similar for both methods (Figure 1c,d). Thus, DeepNull and Baseline produce similar GWAS results when the effect of the covariates on the phenotype is linear.

**Figure 1:**
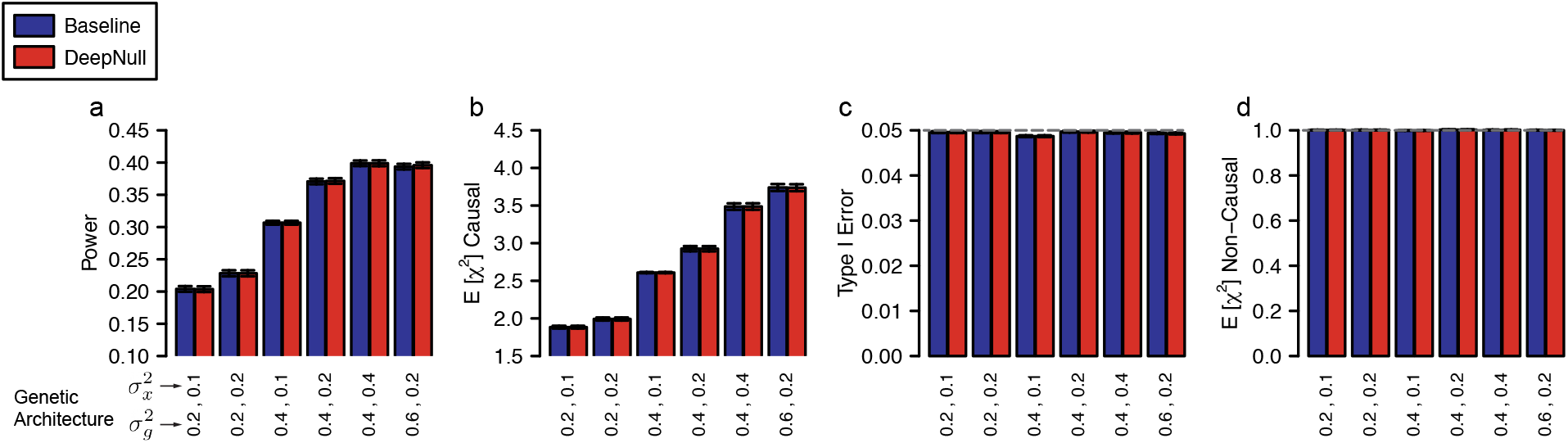
DeepNull and Baseline model achieve similar results under simulated linear covariate effects. (a) Statistical power, (b) Expected Chi-square statistics for variants in the causal chromosome (`chr22`), (c) Type I Error, and (d) Expected Chi-square statistics for variants in the non-causal chromosomes (`chr1` and `chr2`.). In the case of power and the expected Chi-square statistics in the causal chromosome, higher is better. Methods should have a type I error of 0.05 (grey dashed horizontal line). The expected Chi-square statistics for the non-causal chromosomes should be 1 (grey dashed horizontal line). X-axis values indicate the proportion of phenotypic variance explained by genotypes and covariates, respectively. Error bars are the standard error mean for each estimate. None of the quantities shown is significantly different between Baseline and DeepNull (Wilcoxon signed-rank test).

### 2.3 DeepNull increases power in the presence of covariate interactions

We simulated phenotypes using a similar process as described above and used standardized `age`, `sex`, `genotyping_array`, `age`^2^, `age` × `sex`, and `age` × `genotyping_array` as true covariates. However, both DeepNull and Baseline are only given `age`, `sex`, `genotyping_array` as known covariates. This simulation setting explores the case where the true covariates are known but their possible interactions are not. DeepNull has higher statistical power (2%–13% relative improvement) and expected Chi-square statistics for causal variants (2%–20% relative improvement) across all genetic architectures (Figure 2a,b and Table S3). Importantly, both DeepNull and Baseline control type I error and generate similar expected Chi-square statistics for non-causal variants (Figure 2c,d).

**Figure 2:**
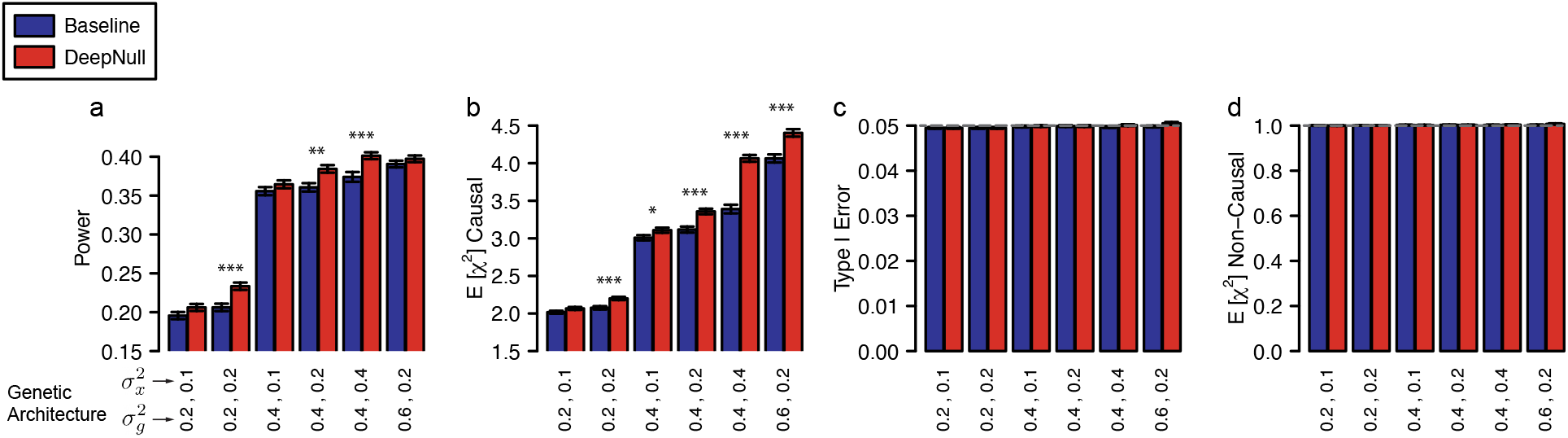
DeepNull increases power with simulated covariate interactions. (a) Statistical power, (b) expected Chi-square statistics for variants in the causal chromosome (`chr22`), (c) Type I Error, and (d) expected Chi-square statistics for variants in the non-causal chromosomes (`chr1` and `chr2`.). In the case of power and expected Chi-square statistics in the causal chromosome, higher is better. Methods should have a type I error of 0.05 which is shown in grey dashed horizontal line. For the non-causal chromosomes, the expected Chi-square statistics should be 1, which is also shown in grey dashed horizontal line. X-axis values indicate the proportion of phenotypic variance explained by genotypes and covariates, respectively. Error bars are the standard error mean for each estimate. The numerical results are shown in Table S3. Indications of *P* value (Wilcoxon signed-rank test) ranges: ^*^*P* ≤ 0.05, ^**^*P* ≤ 0.01, ^***^*P* ≤ 0.001.

### 2.4 DeepNull increases power under non-linear model

We simulated phenotypes using a similar process as described above and again used `age`, `sex`, `genotyping_array`, `age`^2^, `age` × `sex`, and `age` × `genotyping_array` as true covariates. However, here we fix the genetic architecture (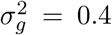 and 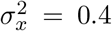) and consider non-linear effects of the covariates on the phenotype by using different non-linear functions for *f* (·) in Equation (10): sin(*x*), exp(*x*), log(|*x*|), and sigmoid(*x*). Again, both DeepNull and Baseline are only given `age`, `sex`, and `genotyping_array` as known covariates. In all cases, DeepNull outperforms Baseline both in terms of statistical power (3%–9% relative improvement) and the expected Chi-square statistics (13%–22% relative improvement) while both methods control type I error (Table S4).

We observed that DeepNull is computationally efficient (see Supplementary Notes) and DeepNull power increases as sample size increases (see Supplementary Notes; Figure S2, Table S5). Finally, DeepNull results are not affected by random seed initialization (see Supplementary Notes; Figure S3).

### 2.5 DeepNull detects more hits for UKB phenotypes than traditional GWAS

To explore whether applying DeepNull is beneficial in non-simulated data, we performed GWAS for ten phenotypes in the UK Biobank using both Baseline and DeepNull: Alkaline phosphatase (ALP), Alanine Aminotransferase (ALT), Aspartate aminotransferase (AST), Apolipoprotein B (ApoB), Calcium, Glaucoma referral probability (GRP), LDL cholesterol (LDL), Phosphate, Sex hormone-binding globulin (SHBG), and Triglycerides (TG), each of which has evidence of potential non-linear relationships between covariates and phenotype (Figures S4–S13). In the case of ALP, ALT, AST, ApoB, Calcium, LDL, SHBG, Phosphate, and TG, the raw phenotypic values are used. `age`, `sex`, and `genotyping_array` were considered as input covariates for the DeepNull DNN. We performed GWAS for these phenotypes using `age`, `sex`, `genotyping_array`, and the top 15 PCs as covariates.

For GRP, we followed the same process as Alipanahi *et al*. [1], in which glaucoma referral probability for an individual is computed by a deep learning model applied to color fundus photographs. Similar to Alipanahi *et al*., we are interested in biological signals for glaucoma that are not driven by the vertical cup-to-disc ratio (VCDR). Thus, we performed GWAS on GRP conditional on VCDR. For the DeepNull DNN, we used `VCDR`, `age`, `sex`, and `genotyping_array` to predict GRP. We then performed GWAS for GRP using `age`, `sex`, `genotyping_array`, the top 15 PCs, `VCDR visit`, `refractive error`, and `image gradability` as covariates.

For all GWAS, we computed the stratified LD score regression (S-LDSC) [3, 13] intercept to determine whether the results were inflated by confounding. We observed that for all ten phenotypes the S-LDSC intercept does not differ significantly from 1, providing no evidence of confounding in our analysis (Table S6). In addition, the SNP-heritability of all phenotypes for both DeepNull and Baseline were estimated from S-LDSC, which showed nominally (but not statistically significantly) larger heritability for DeepNull in all phenotypes except GRP (Table S6).

We observed that DeepNull detects more genome-wide significant hits (i.e. independent lead variants) and loci (independent regions after merging hits within 250 Kbp together; see Methods) than Baseline for all phenotypes examined (Table 1). For example, we find 41% more significant loci (38 vs 26) for GRP using DeepNull compared to the Baseline model. Similarly, in the case of LDL, we detected 202 significant loci using DeepNull compared to the 193 significant loci detected in Baseline (4.6% more hits and 4.7% more loci). In addition, 99 of the DeepNull loci were replicated in the GWAS catalog compared with 96 loci for Baseline (Figure S14). In the case of ApoB, we observed that DeepNull detects 1219 hits compared to 1172 hits detected by Baseline (4.0% improvement) and DeepNull detects 217 significant loci compared to 200 significant loci obtained from Baseline (8.5% improvement; see Table 1). In addition, 166 of the DeepNull loci were replicated in the GWAS catalog compared with 165 loci for Baseline (Figure S15). Overall, the average percentage improvement for DeepNull in significant hits and loci across the ten phenotypes was 6.29% and 7.86%, respectively (Table 1), and the average number of hits and loci improved by 3.29% (851.6 for DeepNull vs 824.4 for Baseline) and 3.93% (222.5 for DeepNull vs 214.1 for Baseline), respectively.

**Table 1:** DeepNull improves association results over the Baseline model on ten phenotypes in UK Biobank. *n* is the sample size, Hits: is the number of independent genome-wide significant hits detected, and Loci is the number of independent regions after merging hits within 250 kb. We use the following abbreviations: ALP (Alkaline phosphatase), ALT (Alanine Aminotransferase), AST (Aspartate aminotransferase), ApoB (Apolipoprotein B), GRP (Glaucoma referral probability), LDL (Low-density lipoprotein), SHBG (Sex hormone-binding globulin), and TG (Triglycerides).

| Pheno | $n$ | #Hits | | %Improve | #Loci | | %Improve |
| --- | --- | --- | --- | --- | --- | --- | --- |
|  |  | Baseline | DeepNull |  | Baseline | DeepNull |  |
| ALP | 416,232 | 1697 | **1759** | 3.65% | 336 | **350** | 4.17% |
| ALT | 416,057 | 371 | **379** | 2.16% | 173 | **174** | 0.58% |
| AST | 414,743 | 337 | **351** | 4.15% | 137 | **145** | 5.84% |
| ApoB | 414,639 | 1172 | **1219** | 4.01% | 200 | **217** | 8.50% |
| Calcium | 381,934 | 726 | **739** | 1.79% | 272 | **281** | 3.31% |
| GRP | 65,896 | 28 | **38** | 35.71% | 26 | **38** | 46.15% |
| LDL | 415,892 | 950 | **993** | 4.53% | 193 | **202** | 4.66% |
| Phosphate | 381,362 | 658 | **664** | 0.91% | 224 | **229** | 2.23% |
| SHBG | 378,459 | 1084 | **1120** | 3.32% | 319 | **323** | 1.25% |
| TG | 416,295 | 1221 | **1254** | 2.70% | 261 | **266** | 1.92% |
| Avg. | 370,151 | 824.4 | **851.6** | 6.29% | 214.1 | **222.5** | 7.86% |

To further understand the source of the DeepNull improvements, we evaluated two additional Baseline models of increasing complexity. The first model, which we call “Baseline+ReLU”, featurizes `age` into five additional covariates by applying the ReLU function at different thresholds (and solely for GRP, also featurizes `VCDR` in the same way). We observed that while Baseline+ReLU generally identified more significant hits and loci than Baseline, DeepNull consistently outperformed both baseline methods (Table S7). The second model, which we call “Second-order Baseline”, extends the Baseline model to include all second-order interactions between `age`, `sex`, and `genotyping_array`: `age`^2^, `age` × `sex`, `age` × `genotyping_array`, and `sex` × `genotyping_array`. Although the additional second-order interaction covariates consistently improve over the Baseline model results, DeepNull detects as many or more significant loci than Second-order Baseline for nine of the 10 phenotypes (Table S8). For AST, LDL, Phosphate, and TG, Second-order Baseline and DeepNull detected similar hits and loci (Tables S9 and S10), providing evidence that the hits and loci not found by the Baseline model, which does not include second-order interactions, are true signals. The utility of DeepNull arises because the optimal order of covariate interactions is not known *a priori* (Table S1) and exhaustively enumerating higher-order interactions may introduce collinearity.

### 2.6 DeepNull improves phenotype prediction for UKB phenotypes

An important advantage of DeepNull is that it provides additional signal for phenotype prediction. Typically, phenotype prediction models are created using a linear combination of common covariates (such as age and sex) and a polygenic risk score (PRS) defined using GWAS association results. Covariate interactions or higher order terms are occasionally included, but typically in an ad-hoc fashion. DeepNull provides a way to easily include potential covariate interactions or higher order terms. When compared to a baseline model that uses `age`, `sex`, and PRS_baseline_ to predict the phenotype, a model that uses `age`, `sex`, PRS_DeepNull_, and `DeepNull_DNN_prediction` performs significantly better in terms of *R*^2^ where *R* is Pearson’s correlation (Figure 3). We calculated *R*^2^ following previous works [28, 33]. We observed that in the case of GRP, LDL, Calcium, and ApoB, DeepNull improves phenotype prediction by 83.42%, 40.33%, 23.90% and 21.61%, respectively. Overall, DeepNull improves phenotype prediction by 23.72% on average for the ten phenotypes analyzed (average *n*=370K; Table S11). In addition, DeepNull has an average *R*^2^ of 0.1940 compared to Baseline average *R*^2^ of 0.1315 (33.65% improvement; Table S11). To determine whether the improved predictive power stems from more accurate GWAS effect size estimates or inclusion of the DeepNull DNN prediction, we examined predictive performance of a model that uses `age`, `sex`, and PRS_DeepNull_ (“DeepNull PRS”). This model produces slightly higher *R*^2^ compared to Baseline for seven of the ten phenotypes, though the difference is not statistically significant for any phenotype (Table S11). Lastly, we compared phenotype prediction of DeepNull to an extended Baseline model that incorporates second order interactions (additional covariates such as `age`^2^, `age` × `sex`, `age` × `genotyping_array`). The extended Baseline model produces similar *R*^2^ to DeepNull for the blood-based phenotypes, but DeepNull increases phenotype prediction of GRP by 11.81% (compare Tables S11 and S8).

**Figure 3:**
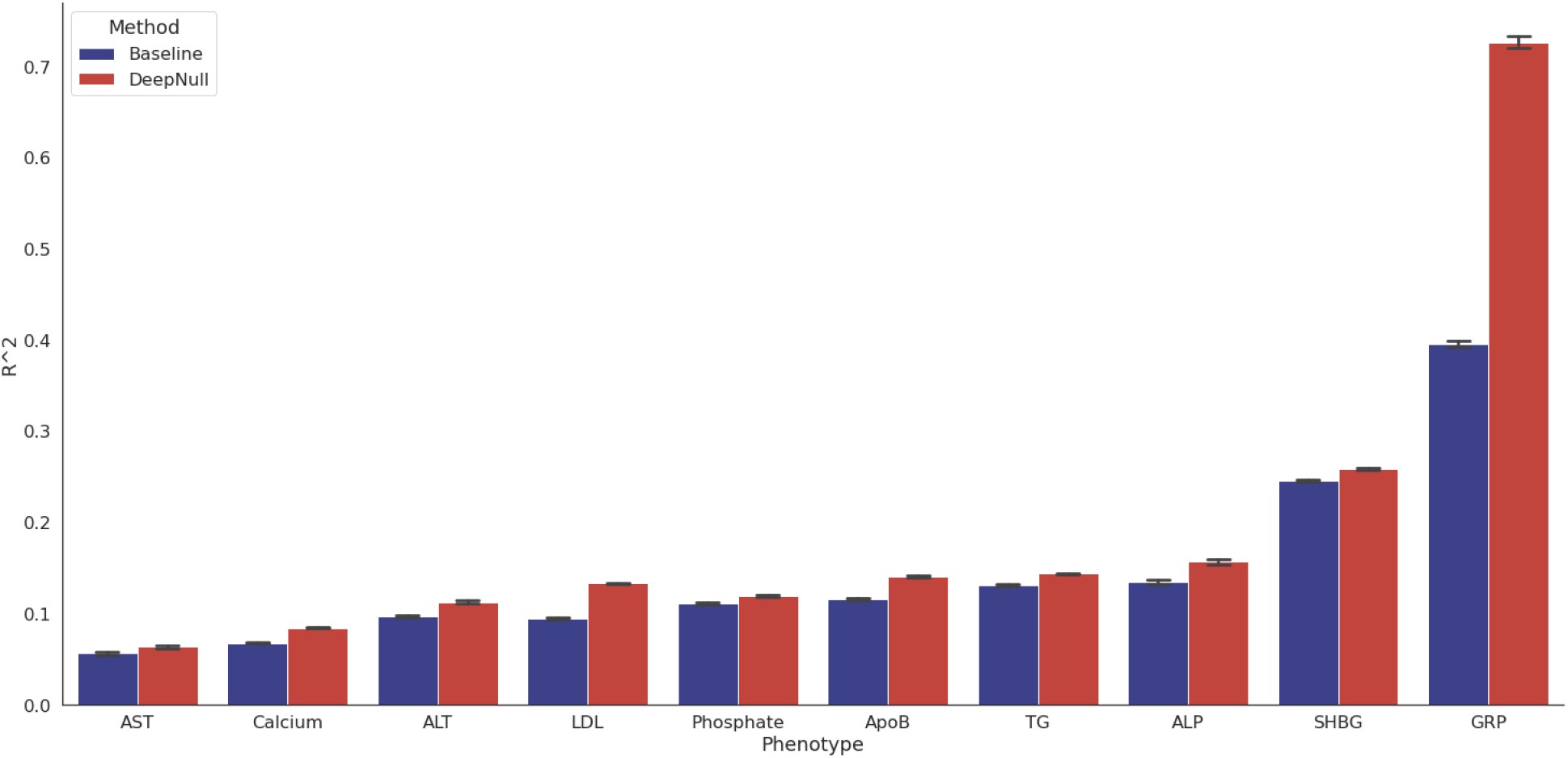
DeepNull improves phenotype prediction compared to Baseline. The X-axis is the phenotype names and the Y-axis is the *R*^2^ where *R* is the Pearson’s correlation between true and predicted value of phenotypes. The error bars indicate the standard error.

### 2.7 Covariates included in DeepNull should remain in the association model

When performing genetic association testing via model shown in Equation (1), the covariates *X* input row-wise to the DNN prediction function *h* are also included as components of the linear term 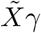. This secondary adjustment for *X* is necessary because *h* captures the association between the covariates *X*_*i*_ and the phenotype *y*_*i*_, but does not capture any association between the covariates *X*_*i*_ and genotype 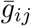. Failure to include *X*_*i*_ in the final association model is comparable to projecting *X*_*i*_ out of *y*_*i*_ but not *g*_*ij*_. To empirically demonstrate the necessity of adjusting *X*_*i*_ in the final association model, we generated phenotypes via

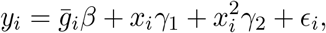

For this simulation only, 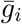 was generated as a continuous random variable, allowing for fine control of the correlation between 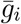 and *x*_*i*_, and the model *h* for predicting *y*_*i*_ from *x*_*i*_ was the oracle model

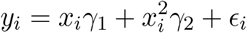

We compare two methods for estimating the genetic effect *β*. The unadjusted model incorporates the prediction *h*(*x*_*i*_) of *y*_*i*_ based on *x*_*i*_ but omits *x*_*i*_ from the association model, emulating the exclusion of covariates provided to DeepNull from the association model as shown in Equation (1),

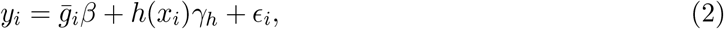

The adjusted model includes both *h*(*x*_*i*_) a linear correction for *x*_*i*_, emulating the application of (1) in practice where the functional form linking *y*_*i*_ and *x*_*i*_ is unknown,

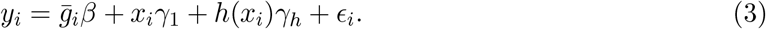

Figure 4 presents the relative bias of the unadjusted and linearly adjusted models for estimating the association parameter *β*. The relative bias for estimating *β* from the generative model, which represents the best possible performance, is also provided. For these simulations *γ*_1_ = 2, *γ*_2_ = −1, and *β* ∈ *{±*1, *±*2, *±*3*}*; the correlation between 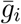 and *x*_*i*_ was 0.5. The unadjusted estimate is generally biased. The magnitude and direction of the bias depend on the coefficients of the generative model. For the unadjusted estimator to be unbiased, 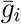 and *x*_*i*_ must be independent. Since the dependence of 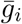 and *x*_*i*_ is seldom clear, the linearly adjusted model is unbiased in either case, and linear adjustment is robust to lower and higher-order covariate influences (Figures S16 and S17), we adopted the fully adjusted model for all other experiments.

**Figure 4:**
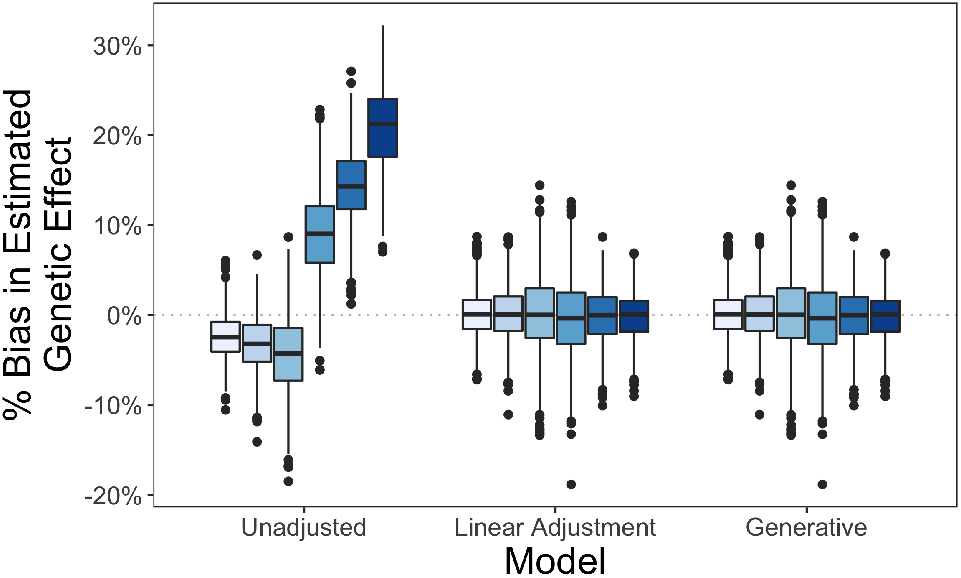
Adjusting for covariates provided to DeepNull during association testing is necessary to avoid bias. The unadjusted model regresses *y*_*i*_ on 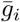 and *h*(*x*_*i*_), the prediction of *y*_*i*_ based on *x*_*i*_, omitting *x*_*i*_ from the association model. This approach results in biased estimation of the genetic effect. The linear adjustment model regresses *y*_*i*_ on 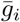, *x*_*i*_, and *h*(*x*_*i*_). This approach is unbiased. The generative model regresses *y*_*i*_ on 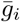, *x*_*i*_, and 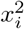. This represents the best possible performance.

## 3 Discussion

A typical GWAS examines the association between genotypes and the phenotype of interest while adjusting for a set of covariates. While covariates potentially have non-linear effects on the phenotype in many real world settings, due to the challenge of specifying the model, GWAS seldom include non-linear terms. Although it is theoretically possible to model the non-linear effects by considering all possible covariate interactions in a linear model, there are multiple limitations: First, the optimal order of covariate interactions is not known *a priori* (Table S1) as it depends on the particular phenotype and set of covariates. Second, adding higher order covariate interactions requires careful analysis to avoid overfitting and collinearity. We proposed a new model, DeepNull, that can model the non-linear effect of covariates on phenotypes when such non-linearity exists. We show that DeepNull can substantially improve phenotype prediction. In addition, we show that DeepNull achieves results similar to a standard GWAS when the covariate effect on the phenotype is linear and can significantly outperform a standard GWAS when the covariate effects are non-linear. DeepNull reduces residual variation, thereby increasing statistical power (Figure S1).

Increasing the statistical power of GWAS is an active area of research that aims to uncover the many variants, each with individually small effect sizes, that collectively explain substantial variation in complex traits. Multiple complementary approaches and methods have been proposed to increase statistical power. The most fundamental one is to increase the sample size of the cohort [44]. However, when resources are limited, the sample size cannot be increased indefinitely, and power can be improved through the use of more refined statistical analyses. Linear mixed models (LMMs) were introduced into GWAS [21, 22, 29–31, 47–50] to allow for the inclusion of related individuals who are not statistically independent. Transformation methods remap or transform the phenotype to make the distribution of phenotypic residuals more nearly normal [14–16, 34, 40]; while normality of the phenotypic residuals is not necessary for valid association testing, standard association tests are most powerful when the residuals are normally distributed. The final class of methods increases power by leveraging external data on the prior biological plausibility of the variants under study. Highly conserved variants, variants in exons, and protein-coding variants all have higher prior probability of being causal than variants in intergenic regions. A series of methods have been developed that use functional data to detect biologically important variants and up-weight their association statistics or reduce their significance thresholds [8, 10, 11, 24, 45, 46]. By modeling non-linear covariate effects on phenotype, DeepNull takes a complementary approach to improving statistical power of GWAS, and thus can be used in combination with some or all of the other approaches discussed above.

We note several limitations of our work. First, while training the DeepNull model, we assume individuals (e.g., samples) are independent. Although this is a general assumption for all machine learning methods and optimization frameworks, this is not necessary true in the presence of related individuals. Thus, we believe that an ML optimizer that can incorporate sample relatedness can speed the training of DeepNull. Importantly, although DeepNull makes the independence assumption during training, this does not mean that type I error is not controlled; DeepNull uses LMM to perform association tests which models the relatedness between individuals. Second, DeepNull does not model possible interactions between genotype-covariate (*G* × *X*) and genotype-genotype (*G* × *G*). This limitation is shared with standard GWAS, and can be overcome only by employing a different statistical model to capture these interactions during association testing.

By accurately modeling non-linear interactions between covariates and the phenotype of interest, DeepNull improves phenotype prediction and association power both in simulations and in ten phenotypes in UKB. Software to perform end-to-end cross-validated training and prediction, and incorporate the results into commonly-used GWAS input data, is freely available (see URLs). The improved performance of DeepNull, combined with its ease of use, suggest that it or similar approaches should be incorporated into the standard framework for performing phenotype prediction and association testing in the future.

## 4 Methods

*Notation*: We use bold capital letters to indicate matrices, non-bold capital letters to indicate vectors, and non-bold lowercase letters to indicate scalars.

### 4.1 Standard GWAS

We consider GWAS of a quantitative trait for a sample of *n* individuals genotyped at *m* SNPs. Let 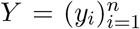 denote the *n* × 1 phenotype vector, where *y*_*i*_ is the phenotypic value of the *i*-th individual, and ***G*** = [*g*_*ij*_] the *n* × *m* sample by SNP genotypes matrix, where *g*_*ij*_ is the minor allele count for the *i*-th individual at the *j*-th variant. Since human genomes are diploid, each variant has 3 possible minor allele counts: *g*_*ij*_ ∈ *{*0, 1, 2*}*. 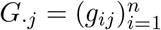 is a vector of minor allele counts for all individuals at the *j*-th SNP. For simplicity, assume the phenotypes and genotypes are standardized to have zero mean and unit variance. Let 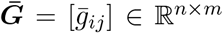 be the standardized version of ***G***, i.e. the empirical mean and variance of 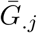 are zero and one, respectively: 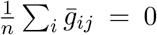 and 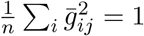 for each *j*-th SNP.

A typical GWAS assumes the effect of each variant on the phenotype is linear and additive. Thus, we have the following generative model:

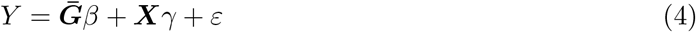

where *β* is the *m* × 1 vector of effect sizes for each variant on the phenotype, ***X*** = [*x*_*ik*_] is the *n* × *q* covariate matrix, including prognostic factors such as age and sex, and *γ* is the *q* × 1 vector of association coefficients for the covariates. For genotypes *g*_*ij*_ ∈ *{*0, 1, 2*}* the assumptions of linearity and additivity are not restrictive. On the other hand, a typical GWAS also assumes that the covariates are linearly associated with the phenotype. This is a far more restrictive assumption if any of the covariates are continuous. 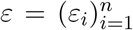 is an *n* × 1 random vector that models the environmental effects and measurement noise.

To perform a GWAS, each variant is individually tested for association with the phenotype. For example, the *j*-th variant is tested for association using the following model:

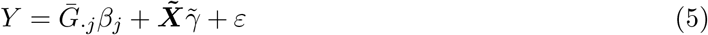

Here 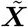 contains the known set of covariates (*e*.*g*. age and sex), in addition to adjustments for confounding that become necessary when the genotypes at SNPs 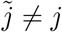 are omitted from the model shown in Equation (4). Confounding due to the presence of genetically related subgroups within the sample, for example subgroups of individuals with common ancestry, is referred to as population structure, and is commonly accounted for by including the top several genetic PCs in 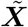 [32, 36, 37].

The model in Equation (5) can be simplified by projecting away the covariates [30, 35]. Define 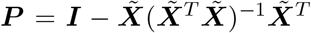, which is the projection onto the orthogonal complement of the linear subspace spanned by ***X***. Multiplying Equation (5) through by ***P*** on the left yields:

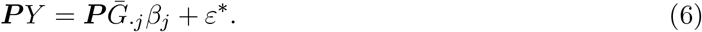

The projected phenotype ***P*** *Y* is the residual from regression of *Y* on 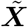. Likewise, 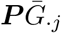 is the residual from regression of 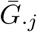 on 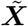. Importantly, if 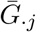 and 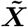 are dependent, which is necessarily true if 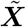 contains confounders of the genotype-phenotype relationship, then 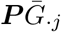 will differ from 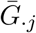. Consequently, an analysis that residualizes only *Y* with respect to 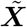 will be misspecified. Instead, to remove dependence on 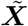, both *Y* and 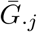 should be residualized in Equation (5).

Though including genotype PCs can control for population structure, it fails to correct for cryptic or family relatedness between individuals [21, 22, 38, 43]. LMMs were introduced to GWAS to overcome these limitations [21, 22, 29–31, 47–50]. LMMs account for random variation in the phenotypic mean that is correlated with the genetic relatedness of the individuals under study, and have proven effective for increasing power even when the kinship among subjects is more distant [30, 31, 48]. We use BOLT-LMM [30, 31] as a framework to perform our analyses and we refer to it as the Baseline method.

### 4.2 DeepNull model

In this work, we consider a model in which the covariates have potentially non-linear effect on the phenotypes. The corresponding generative model for an individual *i* can be written as

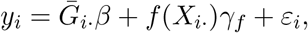

where all variables are defined identically as in formula (4), *f*: ℝ^*q*^ *→*ℝ is any (potentially non-linear) function, 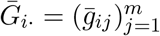, and 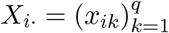. In vector form,

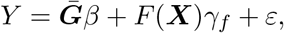

where *F*: ℝ^*n*×*q*^ *→*ℝ^*n*^ is the function that applies *f* to each row of ***X***.

We convert the estimation of *u*_*i*_ = *f* (*X*_*i·*_)*γ*_*f*_ into a learning problem, where we predict *u*_*i*_ using *y*_*i*_ and *X*_*i·*_ as targets and input features, respectively. In other words, we train a model *h* using the covariates *X*_*i·*_ and the phenotype *y*_*i*_ by minimizing

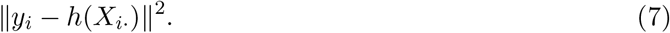

We designed a DNN architecture for modeling the function *h* (Figure 5). It is inspired by residual networks [18] and consists of two components. One component (the shorter path from input to output in Figure 5) is linear, to directly represent the linear effect of the covariates on the phenotype. The other component (the longer path in Figure 5) is a multi-layer perceptron (MLP), to model a potentially non-linear effect of the covariates. The MLP component has 4 hidden layers, all of which use the Rectified Linear Unit (ReLU) activation.

**Figure 5:**
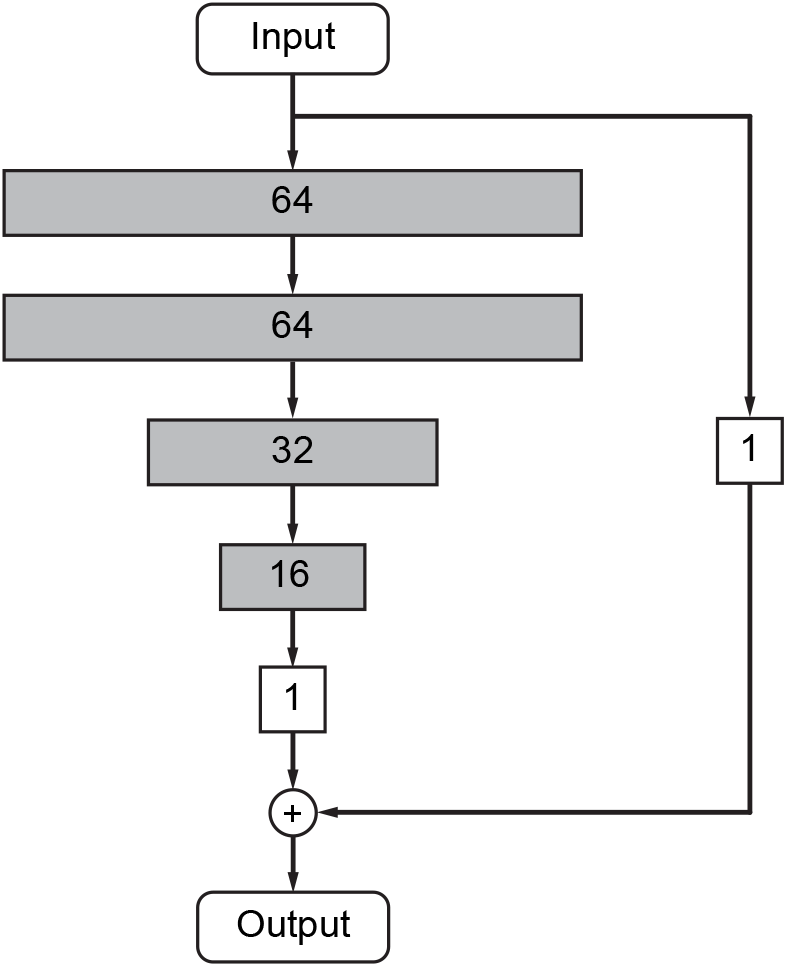
DeepNull DNN model architecture. Each rectangle represents one layer and all layers are fully connected. Shaded layers use the ReLU activation and the non-shaded layers do not use an activation function (i.e. linear connection). The input is the set of known covariates and the output is the predicted phenotype.

In an equation form, the DeepNull model *h* can be written as

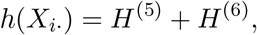

Where

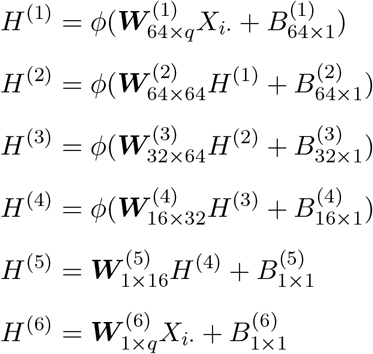

and *ϕ* is the coordinate-wise ReLU function, i.e.

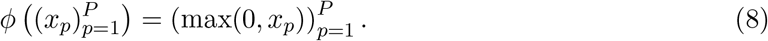

DeepNull learns

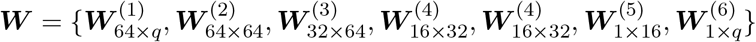

and

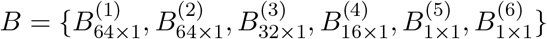

by minimizing the mean squared error (see formula (7)) using the Adam optimizer [25] implemented in Keras for TensorFlow 2. Adam is run with *β*_1_ = 0.9 and *β*_2_ = 0.99. We also used a `batch_size` of 1024 and a `learning_rate` of 10^−4^. We train DeepNull for 1,000 epochs (running DeepNull with more epochs can improve the results with the cost of increasing the training time) without early stopping, batch normalization or dropout. Kernel initializers were set to default (`glorot_uniform`) and bias initializers were set to default (`zeros`).

### 4.3 Performing GWAS using DeepNull

After training DeepNull, we use the following model to test for association between the *j*th variant and the phenotype:

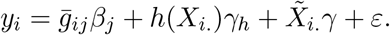

The vectorized form of the above association test is

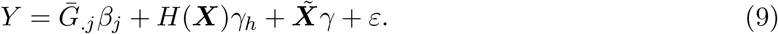

where *H*: ℝ^*n*×*q*^ *→*ℝ^*n*^ is the function that applies *h* to each row of ***X***. Compared to the standard GWAS association model in Equation (5), the DeepNull association model differs only by the inclusion of an extra term *H*(***X***)*γ*_*h*_, where *h*(*X*_*i*._) is the DNN prediction of the phenotype, based on covariates only, and *γ*_*h*_ is a scalar association coefficient. As in the model shown in Equation (5), 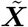 includes both prognostic factors (*e*.*g*. age and sex) and adjustments for confounding (*e*.*g*. genetic PCs) while ***X*** excludes PCs. PCs are excluded as the aim of DeepNull is to predict phenotypes without utilizing genetic data while PCs are computed from genotypes. In addition, higher-order interactions of PCs may capture true biological signals which are not desirable to remove (e.g., conditional associations) in GWAS.

To avoid overfitting, DeepNull should be trained and run on distinct sets of individuals. However, to maximize the GWAS’s statistical power, all individuals in the cohort should receive DeepNull predictions. To satisfy both of these criteria, we split the cohort by individual into *k* partitions. For each selected partition, we train a DeepNull model using data from *k* − 2 of the other partitions and use the remaining partition for validation and model selection. The model that performs best on the validation partition is then used to predict all individuals in the selected partition. The partitioning scheme ensures that each partition is used as the validation/selection partition exactly once.

### 4.4 Simulation framework

We simulate data using the model

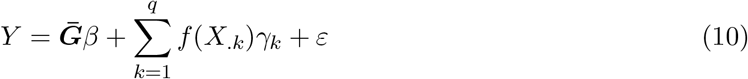

where *X*_.*k*_ is the value of the *k*-th covariate for all individuals, *γ*_*k*_ is the effect size, and *f* (·) is an arbitrary function from ℝ to ℝ, such as the identity function *f* (*x*) = *x* or exponential function *f* (*x*) = exp(*x*). For *j* = 1, …, *m*, the variant effect sizes *β*_*j*_ are drawn independently from a normal distribution with mean zero and variance equal to 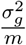 where 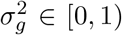 is the proportion of phenotypic variance explained by genotype (i.e., the heritability) and *m* is the number of causal variants: 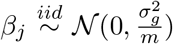. Similarly, the covariate effects are drawn independently from a normal distribution with mean zero and variance equal to 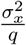 such that 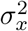 is the proportion of phenotypic variance explained by the covariates: 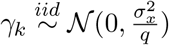. Lastly, *ε* is drawn from another independent normal distribution with mean 0 and variance 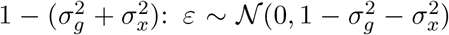. Under this model, 𝔼 (*Y*) = 0 and 𝕍 (*Y*) = 𝔼 (*Y* ^2^) = 1. In the case *f* (·) is the identity function *f* (*x*) = *x*, our simulation framework is similar to previous works [30, 48].

Phenotypes were simulated based on genotypes and covariates from the UKB. `Age`, `sex`, and `genotype_array` were included as covariates. Causal variants were selected uniformly at random from `chr22` such that 1% variants (i.e., 127 variants) were causal. Association testing was performed using BOLT-LMM [31] applied to chromosomes `chr1`, `chr2`, and `chr22`. BOLT-LMM is a linear mixed model that incorporates a Bayesian spike-and-slab prior for the random effects attributed to variants other than that being tested. The prior allows for a non-infinitesimal genetic architecture, in which a mixture of both small and large effect variants influence the phenotype. Specifically, the BOLT-LMM association statistic arises from Equation (9) with the inclusion of an additional random effect 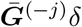. Here 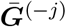 denotes genotype at all variants not on the same chromosome as variant *j*, and the components of *δ* follow the spike-and-slab prior [30].

In our setting, `chr1` and `chr2` are utilized to compute the type I error of the association test, which is the proportion of non-causal variants erroneously associated with the phenotype at a given significance threshold *α* (e.g. *α*=0.05). For null SNPs, the expected *χ*^2^ statistic is 1. Methods that effectively control type I error are compared with respect to their power for correctly rejecting the null hypothesis [9, 12, 41], and their expected *χ*^2^ statistics [30, 31, 48]. Power is defined as the probability of correctly detecting that a variant with a non-zero effect size is causal [9, 12, 41]. Additionally, the expected *χ*^2^ statistic of an association method is a proxy for its prediction accuracy [30, 31, 48].

### 4.5 UKB GWAS evaluation

All GWAS were performed in a subset of UKB individuals of European genetic ancestry. European genetic ancestry was computed as in Alipanahi *et al*. [1]. Briefly, the medioid of the top 15 genetic PC values of all individuals with self-reported “British” ancestry was computed, then the distance from each individual in UKB to the British medioid was computed and all individuals within a distance of 40 were retained. The threshold of 40 was selected based on the 99th percentile of distances of individuals who self-identify as British or Irish.

All GWAS were performed using BOLT-LMM [30, 31] (see URLs) with covariates specific to each experiment. GWAS “hits” were defined as genome-wide significant (i.e. *P* ≤ 5 × 10^−8^) lead variants that are independent at an *R*^2^ threshold of 0.1. Hits were identified using the `--clump` command in PLINK (see URLs). The linkage disequilibrium (LD) calculation was based on a reference panel of 10,000 unrelated subjects of European ancestry from the UKB. The span of each hit was defined based on the set of reference panel variants in LD with the hit at *R*^2^ *≥* 0.1. GWAS “loci” were defined by merging hits within 250 Kbp.

Comparison of two GWAS results *G*_1_ and *G*_2_ for shared and unique hits was performed by examining overlap of the hit spans; a given hit *H*_1_ from *G*_1_ is classified as shared if the span of any hit from *G*_2_ overlaps it, otherwise it is classified as unique.

Comparison of our GWAS with the GWAS catalog (see URLs) was performed analogously to comparing two GWAS. We used `gwas_catalog_v1.0.2-associations_e100_r2021-04-05` and converted coordinates from GRCh38 to GRCh37 using UCSC LiftOver (see URLs) with default parameters. All catalog variants whose “DISEASE/TRAIT” column matched the phenotype of interest and were genome-wide significant were converted into loci by merging variants within 250 Kbp.

## Acknowledgements

This research has been conducted using the UK Biobank Resource application 65275. We are grateful to Alkes L. Price for helpful comments on the manuscript.

## Web Resources

BOLT-LMM software: https://data.broadinstitute.org/alkesgroup/bolt-lmm

BaselineLD annotations: https://data.broadinstitute.org/alkesgroup/ldscore

DeepNull software: github.com/google-health/genomics-research/tree/main/nonlinear-covariate-gwas

GWAS Catalog: https://www.ebi.ac.uk/gwas/

PLINK software: https://www.cog-genomics.org/plink1.9

TensorFlow: https://www.tensorflow.org

UCSC LiftOver: https://genome.ucsc.edu/cgi-bin/hgLiftOver

UK Biobank study: https://www.ukbiobank.ac.uk

