## Supplementary Notes, Table, and Figures for "DeepNull: Modeling non-linear covariate effects improves phenotype prediction and association power"

##### DeepNull is efficient and robust across sample sizes

To assess the computational overhead that DeepNull adds to association testing, we monitored the average time required for the DeepNull DNN to perform one fold of training in a five-fold cross-validation. All experiments were performed on commodity CPU hardware, and in each case DeepNull took 35 min to train. In addition, we observed little difference in DeepNull run times for different $f(\cdot)$ functions.

We varied the sample size (e.g,. $n$) to assess its effect on DeepNull results. We simulated phenotypes for 20K, 50K, 100K, and 200K individuals under the single genetic architecture with $\sigma_g^2 = 0.4$ and $\sigma_x^2 = 0.4$, and considered the two non-linear functions $\exp(x)$ and $\text{sigmoid}(x)$ for $f(\cdot)$ in Equation (10). We observed that both the power and the expected $\chi^2$ statistics for causal variants increase with the sample size (Figure S2) while the type I error is controlled in all cases.

##### DeepNull results are not affected by random seed initialization

It is known that the initial random values assigned to weights at the beginning of training can influence deep learning methods, drastically in some cases. In this section, we investigate whether this phenomenon affects our DeepNull results.

We considered the genetic architecture where we set $\sigma_g^2 = 0.4$ and $\sigma_x^2 = 0.4$ and then simulated phenotypes for 10,000 individuals using UKB genotypes and covariates. We considered the two non-linear functions of $\exp(x)$ and $\text{sigmoid}(x)$ for $f(\cdot)$ in Equation (10). We ran DeepNull prediction using 100 different random seeds for each non-linear function. Then, for each pair of random seeds, we computed the Pearson correlation of the DeepNull prediction for all individuals. As depicted in (Figure S3), those correlations ranged between 0.992 and 1.0. Thus, we conclude that DeepNull results are not significantly affected by the random initial seed.

##### Additional Covariate Adjustment Simulations

*Independent Covariate:* If genotype $\bar{g}_i$ and the covariate $x_i$ are independent, then adjusting for the covariate $x_i$ is not necessary for unbiased estimation of the genetic effect $\beta$. However, adjustment for $x_i$ does improve efficiency, whether by directly including $x_i$ in the association model, or by indirectly including $x_i$ through $h(x_i)$. This is demonstrated in Figure S16. The genotype only model $y_i = \bar{g}_i\beta + \epsilon_i$ is unbiased for estimating $\beta$, but has higher variance than the remaining models. In the absence of association between $\bar{g}_i$ and $x_i$, the unadjusted model (2), the linearly adjusted model (3), and the generative model are all equivalent.

*Cubic Phenotype:* For the quadratic phenotype, the estimate of $\beta$ from the linearly adjusted model

$$y_i = \bar{g}_i\beta + x_i\gamma_1 + h(x_i)\gamma_h + \epsilon_i$$

is in fact numerically identical to that from the generative model

$$y_i = \bar{g}_i\beta + x_i\gamma_1 + x_i^2\gamma_2 + \epsilon_i.$$

To evaluate how the linear adjustment approach performs as the complexity of the relationship between $y_i$ and $x_i$ increases, we repeated the simulation with phenotypes generated from a cubic model

$$y_i = \bar{g}_i\beta + x_i\gamma_1 + x_i^2\gamma_2 + x_i^3\gamma_3 + \epsilon_i.$$

The simulation parameters were $\gamma_1 = 2$, $\gamma_2 = -1$, $\gamma_3 = 1/2$, with correlation 0.5 between $\bar{g}_i$ and $x_i$. As before, the prediction of $y_i$ from $x_i$ was obtained from the oracle model

$$y_i = x_i\gamma_1 + x_i^2\gamma_2 + x_i^3\gamma_3 + \epsilon_i,$$

and the two candidate models for estimating $\beta$ were the unadjusted model (2) and the linearly adjusted model (3).

Results for the cubic simulation (Figure S17) were nearly identical to those from the quadratic simulation (Figure 4). The estimate from the unadjusted model remained biased, and while the estimate from the linearly adjusted model was no longer identical to that from the generative model, the difference was negligible.

### Supplementary Figures

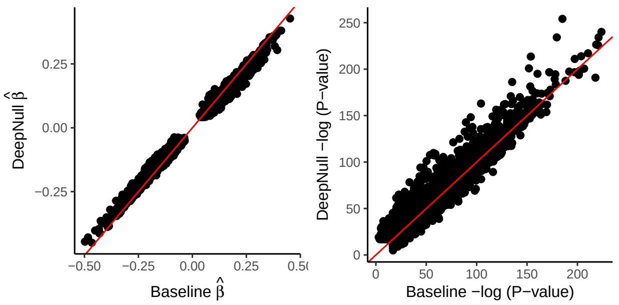


Figure S1: **DeepNull reduces phenotypic residual variation.** a) Estimated effect size where the X-axis is the Baseline and Y-axis is the DeepNull estimated effect size. b) -log of p-value where the X-axis is the Baseline and the Y-axis is the DeepNull significant p-value. Each black dot represents one variant. The red diagonal line indicates the $y = x$. In both panels, we considered all variants that are genome-wide significant by Baseline or DeepNull. We use the simulated setting where $\sigma_g^2 = 0.4$, $\sigma_x^2 = 0.4$, and $f(\cdot) = \exp(x)$.

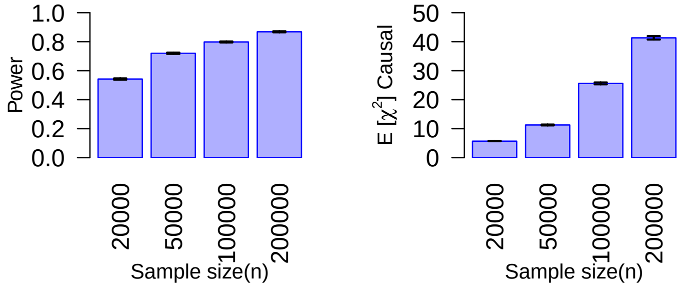


Figure S2: **DeepNull results improve as sample size increases.** a) Y-axis is the statistical power of association. b) Y-axis is the expected $\chi^2$ on the causal chromosome (`chr22`). In both panels, the X-axis is the sample size in the simulated study. The numerical results are shown in Table S5.

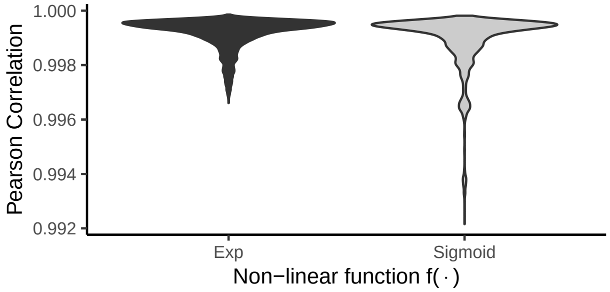


Figure S3: **DeepNull results are not affected by random seed initialization.** Violin plot showing the distribution of Pearson correlations between the DeepNull DNN predictions for pairs of random seed initializers. The narrow range of correlation values shows that the DeepNull DNN is not significantly affected by the choice of random seed. We use $\exp(x)$ and $\text{sigmoid}(x)$ as the non-linear function $f(\cdot)$ from Equation (10).

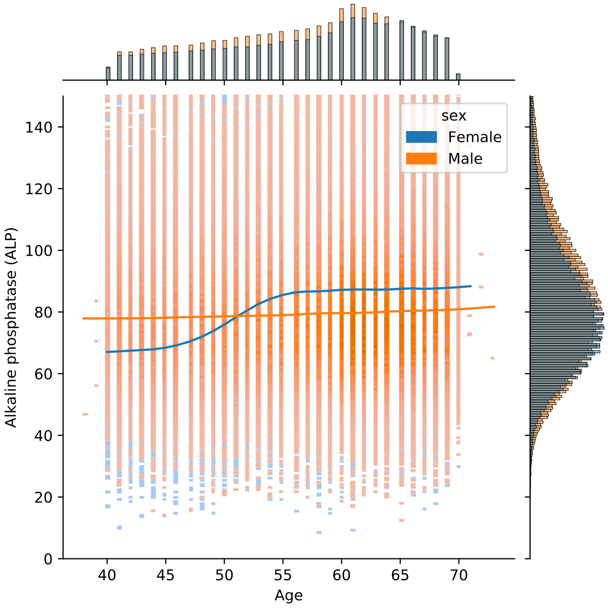


Figure S4: **Alkaline phosphatase (ALP) distribution for each age in UKB.** The blue and orange lines are smoothed non-linear fits of ALP with respect to age.

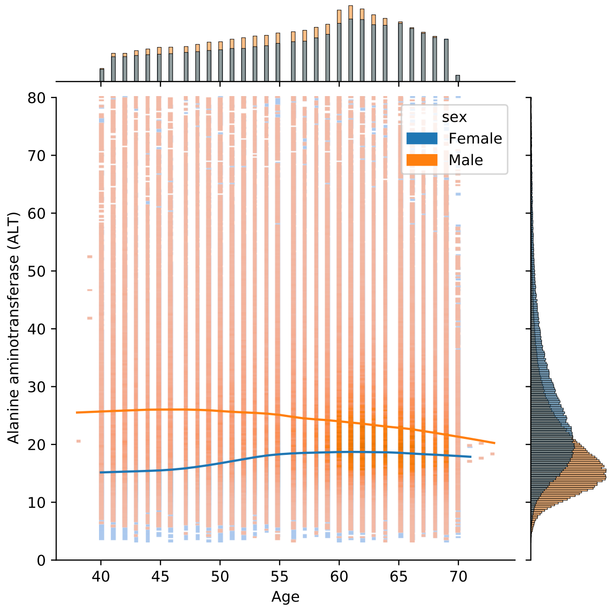


Figure S5: **Alanine aminotransferase (ALT) distribution for each age in UKB.** The blue and orange lines are smoothed non-linear fits of ALT with respect to age.

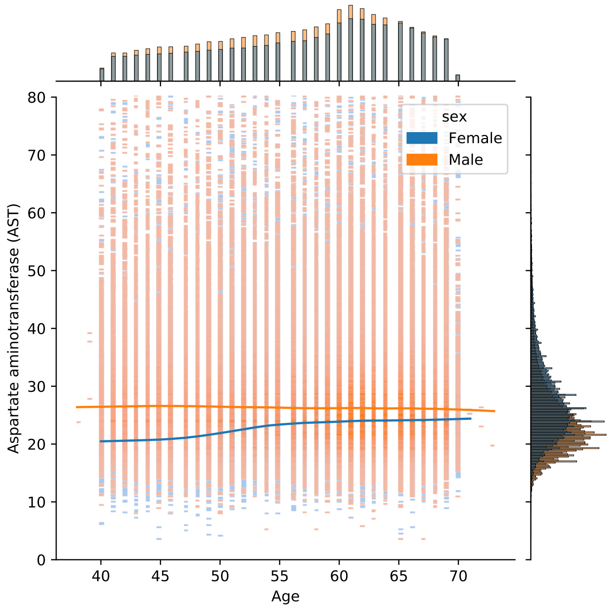


Figure S6: **Aspartate aminotransferase (AST) distribution for each age in UKB.** The blue and orange lines are smoothed non-linear fits of AST with respect to age.

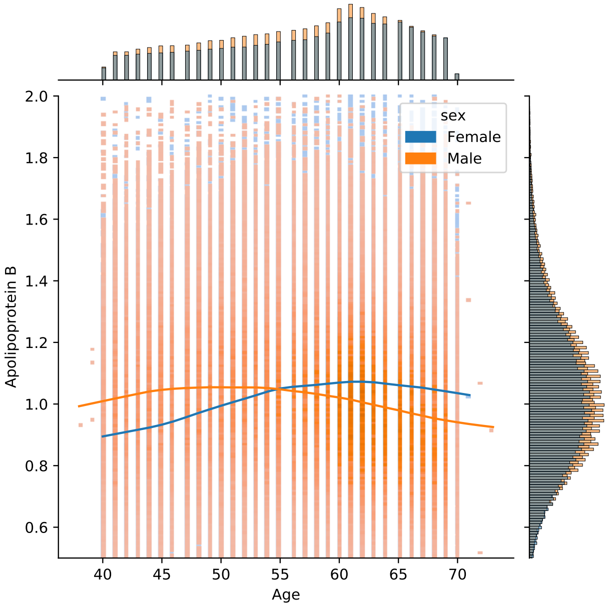


Figure S7: **Apolipoprotein B (ApoB) distribution for each age in UKB.** The blue and orange lines are smoothed non-linear fits of ApoB with respect to age.

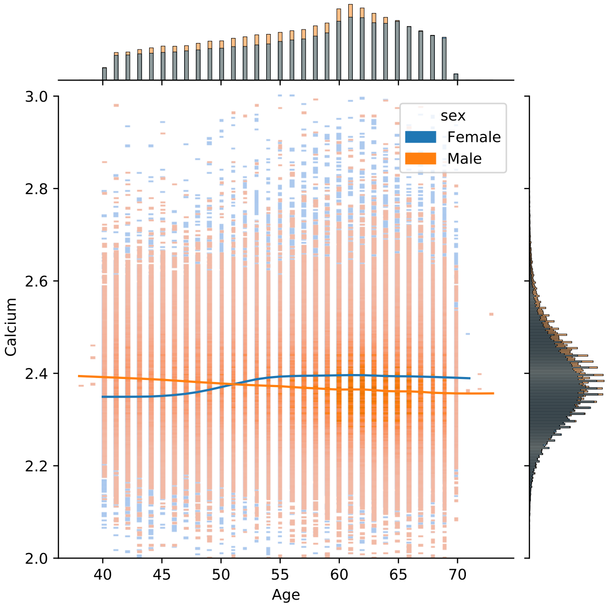


Figure S8: **Calcium distribution for each age in UKB.** The blue and orange lines are smoothed non-linear fits of calcium with respect to age.

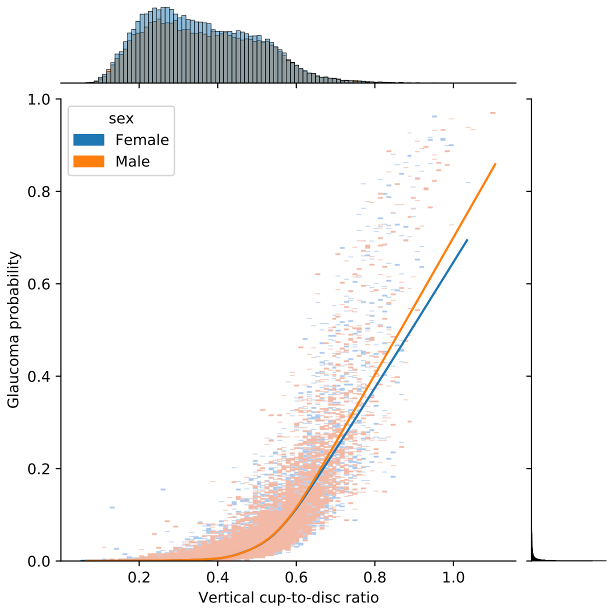


Figure S9: **Glaucoma referral probability (GRP) distribution as a function of vertical cup-to-disc ratio (VCDR) in UKB.** The blue and orange lines are smoothed non-linear fits of GRP with respect to VCDR.

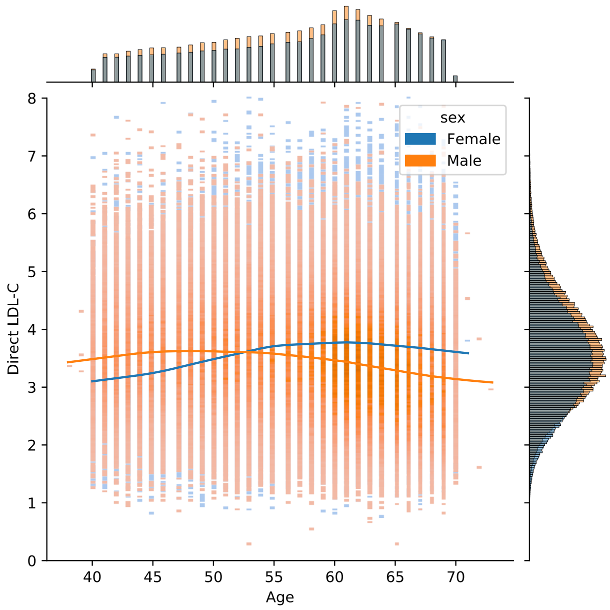


Figure S10: **Low-density lipoprotein (LDL) distribution for each age in UKB.** The blue and orange lines are smoothed non-linear fits of LDL with respect to age.

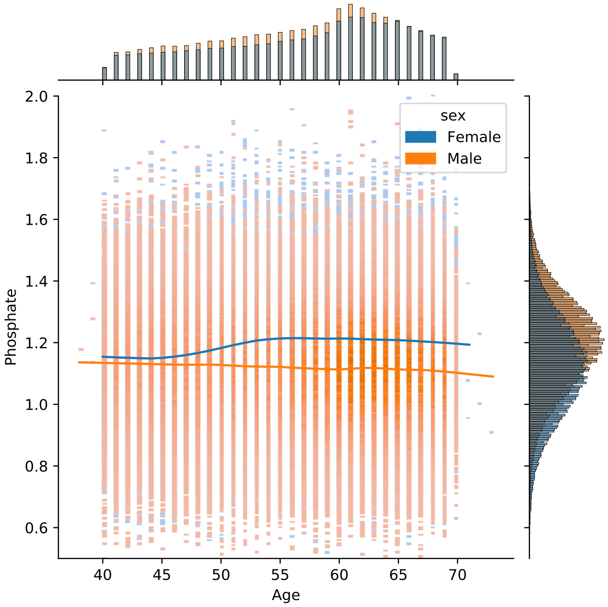


Figure S11: **Phosphate distribution for each age in UKB.** The blue and orange lines are smoothed non-linear fits of Phosphate with respect to age.

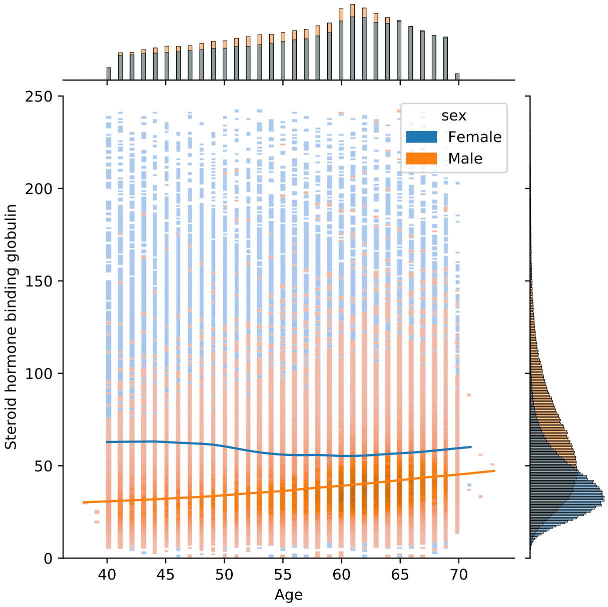


Figure S12: **Sex hormone-binding globulin (SHBG) distribution for each age in UKB.** The blue and orange lines are smoothed non-linear fits of SHBG with respect to age.

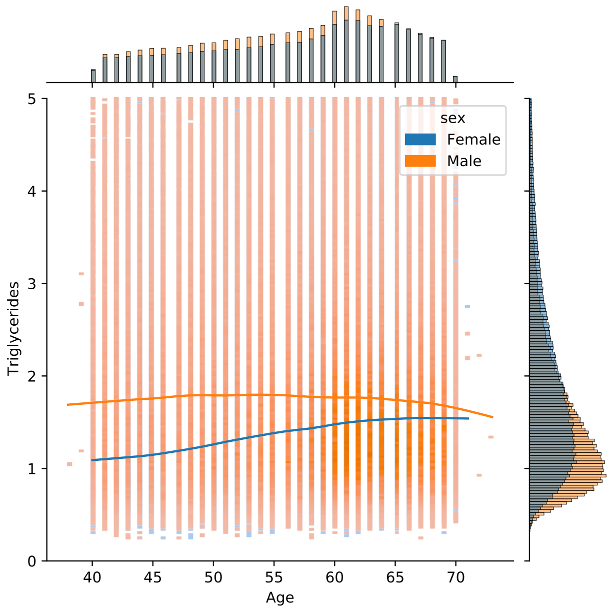


Figure S13: **Triglycerides (TG) distribution for each age in UKB.** The blue and orange lines are smoothed non-linear fits of TG with respect to age.

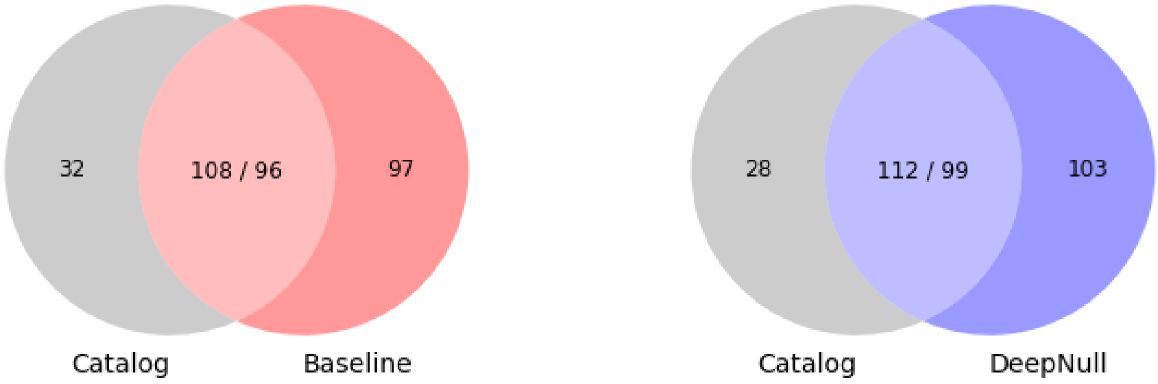


Figure S14: **DeepNull identifies more loci associated with LDL than Baseline.** We computed the overlap of LDL loci for both DeepNull and Baseline with the GWAS catalog trait "LDL cholesterol". Numbers given in the Venn diagrams correspond to locus counts. Because locus overlap is not symmetric (a single GWAS catalog locus may overlap multiple Baseline or DeepNull loci, or vice versa), the shared section of each Venn diagram lists both the number of GWAS catalog loci and the number of Baseline or DeepNull loci.

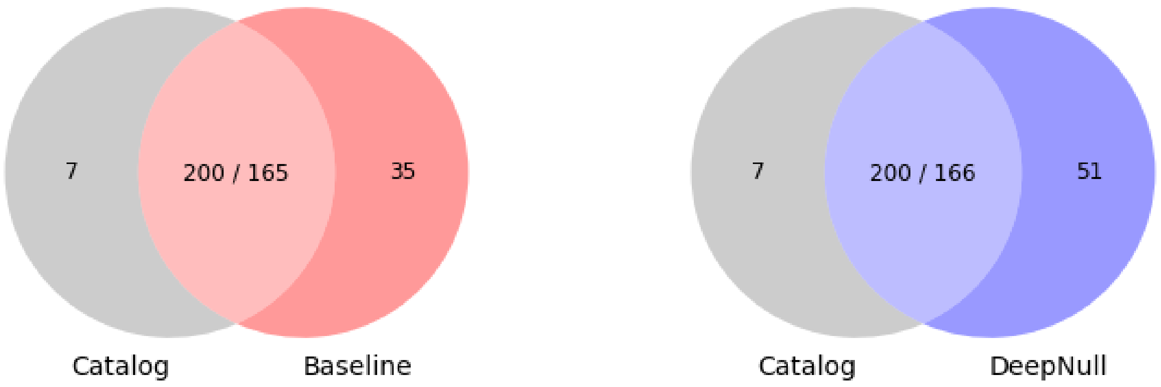


Figure S15: **DeepNull identifies more loci associated with ApoB than Baseline.** We computed the overlap of ApoB loci for both DeepNull and Baseline with the GWAS catalog trait "Apolipoprotein B levels".

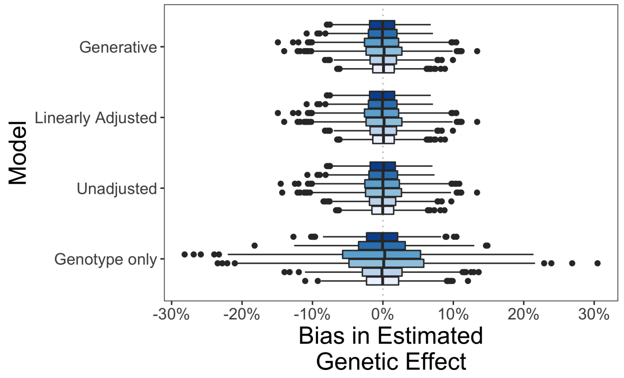


Figure S16: **In the absence of dependence between genotype and the covariate, adjusting for the covariate is more efficient but not necessary for unbiased estimation.** Phenotypes were generated from the quadratic model. The genotype only model regresses $y_i$ on $\bar{g}_i$. The unadjusted model regresses $y_i$ on $\bar{g}_i$ and $h(x_i)$, where $h(x_i)$ is the prediction of $y_i$ based on $x_i$. The linearly adjusted model regresses $y_i$ on $h(x_i)$ and $x_i$. The generative model regresses $y_i$ on $x_i$ and $x_i^2$.

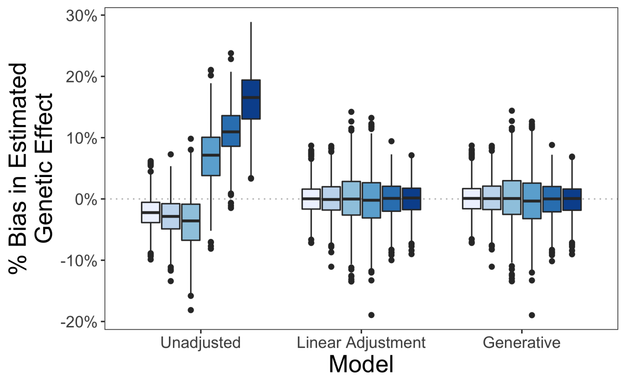


Figure S17: **Linear adjustment provides unbiased estimation of the genetic effect in the case of a cubic phenotype, whereas the unadjusted model remains biased.** The unadjusted model regresses $y_i$ on $\bar{g}_i$ and $h(x_i)$, where $h(x_i)$ is the prediction of $y_i$ based on $x_i$. The linearly adjusted model regresses $y_i$ on $h(x_i)$ and $x_i$. The generative model regresses $y_i$ on $x_i$, $x_i^2$, and $x_i^3$.

### Supplementary Tables

| Pheno | Linear Regression | Linear Regression 2-order interaction | Linear Regression 3-order interaction | Linear Regression 4-order interaction |
|---|---|---|---|---|
| ApoB | 0.0951 | 0.2162 | **0.2237** | 0.1198 |
| BMI | 0.2114 | 0.2414 | 0.2561 | **0.2734** |

Table S1: **The optimal number of covariate interactions for phenotype prediction depends on the phenotype and set of covariates.** We predicted BMI and ApoB phenotypes using `age`, `sex`, `genotyping_array`, 7 diet measures, 4 activity measures, and smoking status, in European individuals in UKB. We used linear regression to perform phenotype prediction with different numbers of covariate interactions. The 7 diet measures used were: 1) `Cooked vegetable intake` (Data-Field: 1289), 2) `Salad/raw vegetable intake` (Data-Field: 1299), 3) `Fresh fruit intake` (Data-Field: 1309), 4) `Bread intake` (Data-Field: 1438), 4) `Coffee intake` (Data-Field: 1498), 5) `Tea intake` (Data-Field: 1488), 6) `Water intake` (Data-Field: 1528), and 7) `Ever added salt to food` (Data-Field: 1478). We used the raw values for the diet measures (excluding the `Ever added salt to food`) and individuals with special values of -1, -3, and -10 were treated as missing. In the case of `Ever added salt to food`, we encoded "Never/rarely" as 0, "prefer not to answer" as missing, and all other values as 1. The 4 activity measures used were: 1) `Summed Metabolic Equivalent Task (MET) minutes per week for all activity` (Data-Field: 22040), 2) `Summed days activity` (Data-Field: 22033), 3) `Summed minutes for walking` (Data-Field: 22034), and 4) `Number of days/week of vigorous physical activity 10+ minutes` (Data-Field: 904). Table cells contain the Pearson correlation ($R$) between predicted and true phenotypic values.

| $\sigma_g^2$ | $\sigma_x^2$ | Method | Power | $\mathbb{E}[\chi]^2$ Causal chr | Type I Error | $\mathbb{E}[\chi^2]$ Non-causal chr |
|---|---|---|---|---|---|---|
| 0.2 | 0.1 | DeepNull | 0.2037 (0.0044) | 1.8837 (0.0184) | 0.0496 (0.0002) | 1.0012 (0.0015) |
| | | Baseline | 0.2040 (0.0044) | 1.8839 (0.0184) | 0.0496 (0.0002) | 1.0012 (0.0015) |
| 0.2 | 0.2 | DeepNull | 0.2283 (0.0047) | 1.9920 (0.0203) | 0.0496 (0.0002) | 1.0014 (0.0015) |
| | | Baseline | 0.2283 (0.0047) | 1.9922 (0.0203) | 0.0496 (0.0002) | 1.0015 (0.0015) |
| 0.4 | 0.1 | DeepNull | 0.3067 (0.0029) | 2.6091 (0.0081) | 0.0487 (0.0002) | 0.9992 (0.0012) |
| | | Baseline | 0.3067 (0.0029) | 2.6087 (0.0081) | 0.0487 (0.0002) | 0.9992 (0.0012) |
| 0.4 | 0.2 | DeepNull | 0.3715 (0.0043) | 2.9261 (0.0345) | 0.0497 (0.0002) | 1.0028 (0.0016) |
| | | Baseline | 0.3707 (0.0043) | 2.9268 (0.0345) | 0.0497 (0.0002) | 1.0029 (0.0016) |
| 0.4 | 0.4 | DeepNull | 0.3989 (0.0045) | 3.4873 (0.0430) | 0.0495 (0.0002) | 1.0020 (0.0017) |
| | | Baseline | 0.3989 (0.0043) | 3.4866 (0.0430) | 0.0495 (0.0002) | 1.0016 (0.0017) |
| 0.6 | 0.2 | DeepNull | 0.3959 (0.0044) | 3.7374 (0.0461) | 0.0493 (0.0002) | 1.0004 (0.0017) |
| | | Baseline | 0.3937 (0.0043) | 3.7396 (0.0462) | 0.0494 (0.0002) | 1.0008 (0.0017) |

Table S2: **Comparison of Baseline and DeepNull models with only linear effects of covariates to phenotype.** Power is computed as the probability of detecting a variant as causal when the true simulated effect size is non-zero. $\mathbb{E}[\chi^2]$ is the expected chi-square statistics for all variants. All values are computed by averaging over 100 simulated datasets and the values in parentheses are the standard error mean (s.e.m.) for each estimate. $\sigma_g^2$ is the phenotypic variance explained by genetic data (i.e. heritability). $\sigma_x^2$ is the phenotypic variance explained by covariates.

| $\sigma_g^2$ | $\sigma_x^2$ | Method | Power | $\mathbb{E}[\chi^2]$ Causal chr | Type I Error | $\mathbb{E}[\chi^2]$ Non-causal chr |
|---|---|---|---|---|---|---|
| 0.2 | 0.1 | DeepNull | **0.2061 (0.0045)** | **2.0671 (0.0195)** | 0.0495 (0.0002) | 1.0004 (0.0015) |
| | | Baseline | 0.1956 (0.0046) | 2.0169 (0.0195) | 0.0495 (0.0002) | 1.0007 (0.0015) |
| | | Relative % | 5.64 | 2.52 | | |
| 0.2 | 0.2 | DeepNull | **0.2334 (0.0047)** | **2.2000 (0.0215)** | 0.0495 (0.0002) | 1.0006 (0.001) |
| | | Baseline | 0.2062 (0.0048) | 2.0755 (0.0224) | 0.0495 (0.0002) | 1.0010 (0.001) |
| | | Relative % | 13.1 | 6.02 | | |
| 0.4 | 0.1 | DeepNull | **0.3645 (0.0051)** | **3.1081 (0.0339)** | 0.0499 (0.0002) | 1.0023 (0.0016) |
| | | Baseline | 0.3557 (0.0052) | 3.0078 (0.0347) | 0.0498 (0.0002) | 1.0027 (0.0016) |
| | | Relative % | 2.47 | 3.33 | | |
| 0.4 | 0.2 | DeepNull | **0.3843 (0.0049)** | **3.3576 (0.0371)** | 0.0499 (0.0002) | 1.0027 (0.0016) |
| | | Baseline | 0.3606 (0.0054) | 3.1165 (0.0403) | 0.0499 (0.0002) | 1.0031 (0.0017) |
| | | Relative % | 6.7 | 7.9 | | |
| 0.4 | 0.4 | DeepNull | **0.4013 (0.0044)** | **4.0642 (0.0450)** | 0.0501 (0.0002) | 1.0038 (0.0018) |
| | | Baseline | 0.3740 (0.0063) | 3.3896 (0.0574) | 0.0497 (0.0002) | 1.0028 (0.0017) |
| | | Relative % | 7.29 | 19.90 | | |
| 0.6 | 0.2 | DeepNull | 0.3973 (0.0043) | **4.4031 (0.0499)** | 0.0505 (0.0003) | 1.0072 (0.0027) |
| | | Baseline | 0.3906 (0.0043) | 4.0631 (0.0536) | 0.0498 (0.0002) | 1.0031 (0.0018) |
| | | Relative % | - | 8.36 | | |

Table S3: **Comparison of Baseline and DeepNull models with covariate interactions.** This result is obtained from a similar process as Figure 2. All values are computed by averaging over 100 simulated datasets and the values in parentheses are the standard error mean (s.e.m.) for each estimate.

| $f(\cdot)$ | Method | Power | $\mathbb{E}[\chi^2]$ Causal chr | Type I Error | $\mathbb{E}[\chi^2]$ Non-causal chr |
|---|---|---|---|---|---|
| | DeepNull | **0.4028 (0.0048)** | **4.0520 (0.0451)** | 0.0500 (0.0002) | 1.0030 (0.0018) |
| $\sin(x)$ | Baseline | 0.3893 (0.0058) | 3.5788 (0.0540) | 0.0498 (0.0002) | 1.0041 (0.0018) |
| | Relative % | 3.45 | 13.22 | | |
| | DeepNull | **0.4044 (0.0046)** | **4.0026 (0.0443)** | 0.0500 (0.0002) | 1.0024 (0.0017) |
| $\sin(20x)$ | Baseline | 0.3756 (0.0061) | 3.3506 (0.0551) | 0.0501 (0.0002) | 1.0042 (0.0016) |
| | Relative % | 7.73 | 19.42 | | |
| | DeepNull | **0.4022 (0.0041)** | **4.0542 (0.0451)** | 0.0500 (0.0002) | 1.0035 (0.0017) |
| $\exp(x)$ | Baseline | 0.3839 (0.0058) | 3.4504 (0.0575) | 0.0498 (0.0002) | 1.0039 (0.0017) |
| | Relative % | 5.74 | 17.52 | | |
| | DeepNull | **0.4031 (0.0046)** | **4.0324 (0.0455)** | 0.0502 (0.0002) | 1.0040 (0.0019) |
| $\exp(20x)$ | Baseline | 0.3903 (0.0057) | 3.2981 (0.0557) | 0.0500 (0.0002) | 1.0030 (0.0017) |
| | Relative % | 4.35 | 22.2 | | |
| | DeepNull | **0.4002 (0.0043)** | **4.0597 (0.0452)** | 0.0502 (0.0002) | 1.0051 (0.0020) |
| $\log(\lvert x\rvert)$ | Baseline | 0.3720 (0.0065) | 3.488 (0.0679) | 0.0495 (0.0002) | 1.0011 (0.0017) |
| | Relative % | 8.35 | 16.37 | | |
| | DeepNull | **0.4045 (0.0043)** | **4.0598 (0.0453)** | 0.05017 (0.0002) | 1.0046 (0.0018) |
| $\log(\lvert 20x\rvert)$ | Baseline | 0.3720 (0.0065) | 3.4888 (0.0679) | 0.0495 (0.0002) | 1.0011 (0.0017) |
| | Relative % | 8.89 | 16.3 | | |
| | DeepNull | **0.4024 (0.0043)** | **4.0615 (0.0451)** | 0.0501 (0.0002) | 1.003 (0.0017) |
| sigmoid($x$) | Baseline | 0.3751 (0.0062) | 3.3932 (0.0572) | 0.0498 (0.0002) | 1.003 (0.0016) |
| | Relative % | 7.60 | 19.68 | | |
| | DeepNull | **0.4000 (0.0041)** | **4.0526 (0.0450)** | 0.0500 (0.0002) | 1.0031 (0.0018) |
| sigmoid($20x$) | Baseline | 0.3848 (0.0056) | 3.4897 (0.0540) | 0.0500 (0.0002) | 1.0048 (0.0018) |
| | Relative % | 4.16 | 16.13 | | |

Table S4: **Comparison of Baseline and DeepNull models under a simulation of a non-linear effect of the covariates on the phenotype.** We considered the genetic architecture where we set $\sigma_g^2 = 0.4$ and $\sigma_x^2 = 0.4$. This result is obtained from similar process as Table S2 while using `age`, `sex`, `genotyping_array`, `age`$^2$, `age`$\times$`sex`, `age`$\times$`genotyping_array` as true covariates to simulate datasets, however, both Baseline and DeepNull use `age`, `sex`, and `genotype_array` as input covariates. All values are averaged over 100 simulations and the parenthetical values are the standard errors of the mean for each estimate.

| $f(\cdot)$ | $n$ | Power | $\mathbb{E}[\chi^2]$ Causal chr | Type I error | $\mathbb{E}[\chi^2]$ Non-causal chr |
|---|---|---|---|---|---|
| | 20,000 | 0.5425 (0.0052) | 5.7107 (0.0604) | 0.0497 (0.0003) | 0.9999 (0.0024) |
| Exp | 50,000 | 0.7202 (0.0053) | 11.2885 (0.1828) | 0.0543 (0.0011) | 1.0413 (0.0096) |
| | 100,000 | 0.7982 (0.0040) | 25.6248 (0.3505) | 0.0518 (0.0007) | 1.0210 (0.0065) |
| | 200,000 | 0.8683 (0.0042) | 41.3273 (0.6022) | 0.0495 (0.0002) | 1.0013 (0.0019) |
| | 20,000 | 0.5414 (0.0051) | 5.7199 (0.0609) | 0.0497 (0.0003) | 1.0004 (0.0024) |
| Sigmoid | 50,000 | 0.7173 (0.0054) | 11.2115 (0.1757) | 0.0534 (0.0010) | 1.0337 (0.0084 |
| | 100,000 | 0.7976 (0.0039) | 25.5486 (0.3420) | 0.0516 (0.0007) | 1.0179 (0.0060) |
| | 200,000 | 0.8684 (0.0042) | 41.2968 (0.6049) | 0.0494 (0.0002) | 1.0002 (0.0019) |

Table S5: **DeepNull results improve as sample size increases.** We computed the power and expected $\chi^2$ statistics of the causal chromosome, type I error, and the expected $\chi^2$ statistics of the non-causal chromosomes. We considered the genetic architecture where we set $\sigma_g^2 = 0.4$ and $\sigma_x^2 = 0.4$ and then simulated phenotypes for different number of individuals ($n$) using UKB genotype and covariates. We considered two non-linear functions of $\exp(x)$ and sigmoid($x$) for $f(\cdot)$ in Equation (10). All values are computed by averaging over 100 simulated datasets and the values in parentheses are the standard error mean (s.e.m.) for each estimate.

| Pheno | S-LDSC Intercept | | S-LDSC SNP-heritability | |
|---|---|---|---|---|
| | **Baseline** | **DeepNull** | **Baseline** | **DeepNull** |
| ALP | 1.0722 (0.0713) | 1.0751 (0.0733) | 0.1901 (0.0192) | 0.1945 (0.0196) |
| ALS | 1.0440 (0.0222) | 1.0452 (0.0225) | 0.0853 (0.0065) | 0.0865 (0.0066) |
| AST | 1.0345 (0.0194) | 1.0352 (0.0193) | 0.0606 (0.0054) | 0.0610 (0.0054) |
| ApoB | 1.0724 (0.0291) | 1.0762 (0.0298) | 0.1277 (0.0131) | 0.1312 (0.0131) |
| Calcium | 1.0962 (0.0309) | 1.0979 (0.0315) | 0.1260 (0.0077) | 0.1281 (0.0079) |
| GRP | 0.9991 (0.0086) | 1.0044 (0.0089) | 0.0237 (0.0113) | 0.0221 (0.0123) |
| LDL | 1.0540 (0.0262) | 1.0575 (0.0267) | 0.1120 (0.0109) | 0.1161 (0.0113) |
| Phosphate | 1.0423 (0.0242) | 1.0432 (0.0244) | 0.1312 (0.0088) | 0.1327 (0.0089) |
| SHBG | 1.1677 (0.0799) | 1.1724 (0.0807) | 0.1533 (0.0138) | 0.1559 (0.0139) |
| TG | 1.1133 (0.04) | 1.1158 (0.0406) | 0.1553 (0.0127) | 0.1572 (0.0129) |

Table S6: **S-LDSC results on Baseline and DeepNull.** We computed the S-LDSC intercept and SNP-heritability. In all phenotypes the S-LDSC intercept is the same for both Baseline and DeepNull which indicates no confounding. Values in parentheses are the standard error mean (s.e.m) obtained from S-LDSC.

| Pheno | #Hits | | | #Loci | | |
|---|---|---|---|---|---|---|
| | **Baseline** | **Baseline + ReLU** | **DeepNull** | **Baseline** | **Baseline + ReLU** | **DeepNull** |
| ALP | 1697 | 1705 | **1759** | 336 | 335 | **350** |
| ALT | 371 | 375 | **379** | 173 | 172 | **174** |
| AST | 337 | 341 | **351** | 137 | 140 | **145** |
| ApoB | 1172 | 1192 | **1219** | 200 | 209 | **217** |
| Calcium | 726 | 732 | **739** | 272 | 275 | **281** |
| GRP | 28 | **38** | **38** | 26 | 36 | **38** |
| LDL | 950 | 956 | **993** | 193 | 198 | **202** |
| Phosphate | 658 | **667** | 664 | 224 | **230** | 229 |
| SHBG | 1084 | 1081 | **1120** | 319 | 316 | **323** |
| TG | 1221 | 1229 | **1254** | 261 | 264 | **266** |

Table S7: **DeepNull consistently detects more hits and loci compared to Baseline and Baseline+ReLU.** Baseline+ReLU is a model where we add additional covariates to Baseline model. We obtained these additional covariates by applying the ReLU function on the covariate of interest using 5 different thresholds. For all phenotypes excluding the GRP, we add 5 additional ReLU based on age. In the case of GRP, we used 10 additional ReLU where 5 covariates are computed using age and another 5 are computed using VCDR.

| Pheno | #Hits | #Loci | $R^2$ | S-LDSC Intercept | S-LDSC SNP-heritability |
|---|---|---|---|---|---|
| ALP | 1750 | 343 | 0.1509 (0.1464, 0.1560) | 1.0730 (0.0727) | 0.1934 (0.0195) |
| ALT | 377 | 173 | 0.1121 (0.1089, 0.1152) | 1.0456 (0.0226) | 0.0864 (0.0066) |
| AST | 351 | 145 | 0.0643 (0.0615, 0.0673) | 1.0352 (0.0193) | 0.0611 (0.0054) |
| ApoB | 1231 | 211 | 0.1491 (0.1470, 0.1508) | 1.0816 (0.0297) | 0.1305 (0.0129) |
| Calcium | 743 | 280 | 0.0882 (0.0862, 0.0897) | 1.0981 (0.0315) | 0.1286 (0.008) |
| GRP | 38 | 35 | 0.6492 (0.6371, 0.6595) | 0.9907 (0.0091) | 0.0372 (0.0124) |
| LDL | 991 | 200 | 0.1337 (0.1318, 0.1358) | 1.0598 (0.0268) | 0.1162 (0.0113) |
| Phosphate | 663 | 226 | 0.1189 (0.1171, 0.1207) | 1.0457 (0.0243) | 0.1318 (0.0089) |
| SHBG | 1113 | 319 | 0.2624 (0.2596, 0.2644) | 1.1738 (0.0809) | 0.1565 (0.0140) |
| TG | 1256 | 267 | 0.1440 (0.1419, 0.1461) | 1.1153 (0.0407) | 0.1575 (0.0129) |

Table S8: **Baseline model with additional second order interaction covariates.** We ran Baseline method with additional covariates that include `age`$^2$, `age` × `sex`, `age` × `genotyping_array`, and `sex` × `genotyping_array`. These covariates supplement the top 15 PCs, `age`, `sex`, and `genotyping_array` included in the Baseline model. $R^2$ is measurement of the phenotype prediction where $R$ is the Pearson correlation between true phenotype and predicted values. The phenotype prediction is the combination of PRS computed using the PLINK `--score` method and linear effects of covariates to phenotype. Here, we show the 95% confidence intervals for the squared Pearson's correlation measuring the correlation between the models predicted value and the true values. We compute the S-LDSC intercept and SNP-heritability. We observe that in all four phenotypes the S-LDSC intercept is the same for both Baseline and DeepNull which indicates no confounding. Values in parentheses for S-LDSC intercept and SNP-heritability are the standard error mean (s.e.m).

| Pheno | Baseline vs DeepNull | | | Second-order vs DeepNull | | |
|---|---|---|---|---|---|---|
| | Baseline-only | Shared | DeepNull-only | Second-only | Shared | DeepNull-only |
| ALP | 8 | 1737 | 22 | 7 | 1749 | 10 |
| ALT | 5 | 376 | 3 | 1 | 375 | 4 |
| AST | 1 | 341 | 10 | 0 | 351 | 0 |
| ApoB | 4 | 1196 | 23 | 4 | 1211 | 8 |
| Calcium | 8 | 719 | 20 | 6 | 733 | 6 |
| GRP | 18 | 8 | 30 | 19 | 16 | 22 |
| LDL | 6 | 976 | 17 | 2 | 989 | 4 |
| Phosphate | 8 | 651 | 13 | 2 | 660 | 4 |
| SHBG | 10 | 1106 | 14 | 5 | 1111 | 9 |
| TG | 6 | 1239 | 15 | 2 | 1252 | 2 |

Table S9: **Number of significant hits shared between DeepNull and Baseline and Second-order Baseline.** In the case of Baseline, we performed GWAS analysis while including `age`, `sex`, `genotyping_array`, and top 15 PCs as possible covariates. In the case of Second-order (2-order) Baseline, we ran Baseline model with additional covariates that includes `age`$^2$, `age` × `sex`, `age` × `genotyping_array`, and `sex` × `genotyping_array`. These covariates supplement the top 15 PCs, `age`, `sex`, and `genotyping_array` included in the Baseline model. "-only" indicates hits that are detected only by one the methods while the "Shared" indicates DeepNull hits that overlap one or more hits from the baseline method. Note that shared hits are not symmetric, but the qualitative results are the same as if we reported the shared baseline method hits.

| Pheno | Baseline vs DeepNull | | | Second-order vs DeepNull | | |
|---|---|---|---|---|---|---|
| | Baseline-only | Shared | DeepNull-only | Second-only | Shared | DeepNull-only |
| ALP | 4 | 333 | 17 | 4 | 340 | 10 |
| ALT | 3 | 171 | 3 | 1 | 172 | 2 |
| AST | 1 | 136 | 9 | 0 | 145 | 0 |
| ApoB | 1 | 199 | 18 | 2 | 210 | 7 |
| Calcium | 8 | 262 | 19 | 6 | 275 | 6 |
| GRP | 18 | 8 | 30 | 19 | 16 | 22 |
| LDL | 5 | 188 | 14 | 2 | 198 | 4 |
| Phosphate | 8 | 217 | 12 | 2 | 225 | 4 |
| SHBG | 7 | 310 | 13 | 5 | 314 | 9 |
| TG | 6 | 254 | 12 | 1 | 265 | 1 |

Table S10: **Number of significant loci shared between DeepNull and Baseline and Second-order Baseline.** This analysis is similar to Table S9; however, here we report the number of loci instead of hits.

| Pheno | Baseline | DeepNull PRS | DeepNull | % Δ (P) |
|---|---|---|---|---|
| ALP | 0.1353 (0.1312, 0.1397) | 0.1363 (0.1323, 0.1406) | 0.1569 (0.1523, 0.1623) | 16.01 ($3.93\times10^{-52}$) |
| ALT | 0.0970 (0.0940, 0.0997) | 0.0970 (0.0940, 0.0997) | 0.1127 (0.1094, 0.1157) | 16.16 ($2.84\times10^{-35}$) |
| AST | 0.0566 (0.0541, 0.0595) | 0.0574 (0.0549, 0.0602) | 0.0642 (0.0616, 0.0672) | 13.32 ($5.07\times10^{-14}$) |
| ApoB | 0.1159 (0.1142, 0.1173) | 0.1166 (0.1149, 0.1181) | 0.1410 (0.1390, 0.1424) | 21.61 ($6.01\times10^{-75}$) |
| Calcium | 0.0682 (0.0666, 0.0697) | 0.0690 (0.0674, 0.0706) | 0.0845 (0.0827, 0.0860) | 23.90 ($1.22\times10^{-44}$) |
| GRP | 0.3958 (0.3889, 0.4022) | 0.3950 (0.3872, 0.4011) | 0.7259 (0.7124, 0.7366) | 83.42 ($\leq1.00\times10^{-200}$) |
| LDL | 0.0950 (0.0935, 0.0967) | 0.0956 (0.0940, 0.0973) | 0.1334 (0.1315, 0.1352) | 40.33 ($6.56\times10^{-189}$) |
| SHBG | 0.2454 (0.2428, 0.2475) | 0.2450 (0.2425, 0.2471) | 0.2581 (0.2555, 0.2602) | 5.19 ($7.74\times10^{-14}$) |
| Phosphate | 0.1111 (0.1093, 0.1129) | 0.1113 (0.1095, 0.1131) | 0.1197 (0.1178, 0.1214) | 7.70 ($2.53\times10^{-10}$) |
| TG | 0.1315 (0.1296, 0.1337) | 0.1319 (0.1299, 0.1341) | 0.1440 (0.1420, 0.1462) | 9.58 ($1.41\times10^{-19}$) |
| Avg. | 0.1451 | 0.1455 | 0.1940 | 23.72 |

Table S11: **DeepNull improves phenotype prediction in terms of Pearson's $R^2$.** We evaluated three models, where each includes `age`, `sex`, and `genotyping_array` as covariates. The "Baseline" model includes a PRS computed using the PLINK `--score` method and linear effect of covariates to phenotype. The "DeepNull PRS" model includes a PRS computed in the same way except using association results from DeepNull, and "DeepNull" is a model that includes both the DeepNull-based PRS and the DeepNull prediction (non-linear effect of covariates to phenotype). Here, we show the 95% confidence intervals for the squared Pearson's correlation between the model-predicted value and the true value. We see that Baseline and "DeepNull PRS" have similar performance with no statistically significant differences. However, the inclusion of the covariate non-linear prediction produces a statistically significant improvement when compared with Baseline or "DeepNull PRS". We compute the overall improvement in two ways: 1) Averaging the relative improvement, which is computed by taking the average of Δ over 10 traits, is 23.72% or 2) Computing the relative improvement over averaged $R^2$ of Baseline and DeepNull. In this case, we have an averaged $R^2$ of 0.1451 for Baseline while DeepNull has an averaged $R^2$ of 0.1940, this indicates a 33.65% improvement.
